# Genetic screening for single-cell variability modulators driving therapy resistance

**DOI:** 10.1101/638809

**Authors:** Eduardo A. Torre, Eri Arai, Sareh Bayatpour, Lauren E. Beck, Benjamin L. Emert, Sydney M. Shaffer, Ian A. Mellis, Mitchell Fane, Gretchen Alicea, Krista A. Budinich, Ashani Weeraratna, Junwei Shi, Arjun Raj

**Affiliations:** Biochemistry and Molecular Biophysics Graduate Group, Perelman School of Medicine, University of Pennsylvania, Philadelphia, PA; Department of Bioengineering, School of Engineering and Applied Science, University of Pennsylvania, Philadelphia, PA; Department of Cancer Biology, Perelman School of Medicine, University of Pennsylvania, Philadelphia, PA; The Wistar Institute, Philadelphia, PA; Department of Pathology and Laboratory Medicine, Perelman School of Medicine, University of Pennsylvania, Philadelphia, PA; Genomics and Computational Biology Graduate Group, Perelman School of Medicine, University of Pennsylvania, Philadelphia, PA; Department of Genetics, Perelman School of Medicine, University of Pennsylvania, Philadelphia, PA; Sidney Kimmel Cancer Center, Johns Hopkins School of Medicine, Baltimore, MD

**Author notes:** Equal contribution (Eri Arai, Sareh Bayatpour).

## Abstract

Cellular plasticity describes the ability of cells to transition from one set of phenotypes to another. In the context of cancer therapeutics, plasticity refers to transient fluctuations in the molecular state of tumor cells, driving the formation of rare cells primed to survive drug treatment and ultimately reprogram into a stably resistant fate. However, the biological processes governing this cellular plasticity remain unknown. We used CRISPR/Cas9 genetic screens to reveal genes that affect cell fate decisions by altering cellular plasticity across a range of functional categories. We found that cellular plasticity and cell fate decision making can be decoupled in that factors can affect cell fate decisions in both plasticity-dependent and independent manners. We discovered a novel mode of altering resistance based on cellular plasticity that, contrary to known mechanisms, pushes cells towards a more differentiated state. We further confirmed our prediction that manipulating cellular plasticity before the addition of the main therapy would result in changes in therapy resistance more than concurrent administration. Together, our results indicate that identifying pathways modulating cellular plasticity has the potential to alter cell fate decisions and may provide a new avenue for treating drug resistance.

## Introduction

Plasticity is often used to describe the ability of cells to transition from one phenotype to another, at times enabling cells to adapt and survive in the face of a variety of stimuli and challenges, for instance, in regeneration, wound healing, and the induction of pluripotency.

Plasticity itself can typically be decomposed into a stimulus-independent and subsequent stimulus-dependent phase. The first phase typically consists of individual, often rare, cells within the population being “primed” for the cell fate transition. Then, upon the stimulus, these primed cells are selectively reprogrammed to adopt the new phenotype. Thus, a major question in single cell biology has been determining the molecular differences specific to these rare primed cells before a stimulus and connecting the molecular profile of these primed cells to their ultimate fate after the stimulus reprograms them. Recently, a number of studies have developed the link between cellular priming and cell fate that underlies plasticity in a number of contexts ^1–8^. However, to date, little is known about the pathways that can manipulate the fluctuations that drive this cellular priming and whether that can affect their subsequent fates, leaving their molecular basis and potential for therapeutic application largely unrealized.

Therapy resistance in melanoma is an excellent example of cellular plasticity ^9, 10^. Therapies such as vemurafenib designed to inhibit particular oncogenic targets can often kill most of the tumor cells, but a few remaining cells can continue to proliferate, ultimately repopulating the tumor. While the mechanisms underlying this therapy resistance can sometimes be the result of a genetic mutation, many recent studies, both in melanoma and other cancers, suggest that cellular plasticity may also dictate which cells are able to survive drug treatment, with rare primed cells being reprogrammed by the addition of drug into a stably resistant state ^8, 11–21^. In melanoma, this rare primed cellular state, which we have also previously referred to as the pre-resistant cellular state, is often marked by transiently high expression of several resistance marker genes, such as *EGFR*, *NGFR* and *AXL* (Fig. 1A, top). Once these cells are exposed to drug, they are reprogrammed into a new cellular fate in which the transient primed phenotype is converted to a stably drug-resistant phenotype characterized by massive changes in signaling and gene expression profiles. This paradigm of resistance has a number of critical differences from the more conventional model of mutational causes of drug resistance—notably, while genetic mutations largely arise through spontaneous, stochastic processes, non-genetic fluctuations that drive the primed cellular states can in principle occur due to the changes in activity of specific biological pathways. Targeting these pathways specifically could have the potential to enhance or inhibit the formation of cells in the primed state independent of the addition of drug. We were thus interested in dissecting the molecular regulators of cellular priming and how those might consequently affect the ultimate drug-resistant fates that cells can adopt.

**Figure 1.**
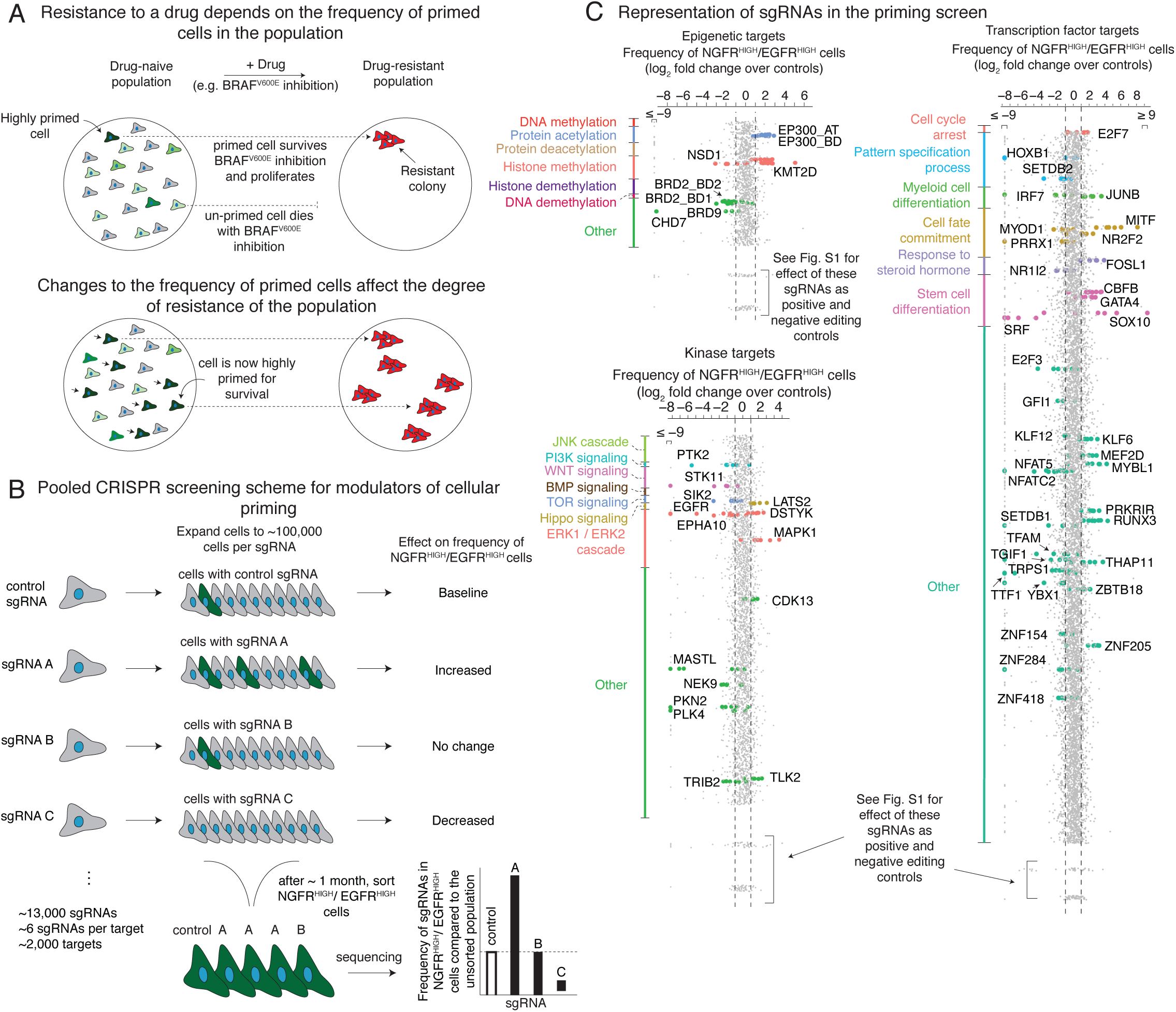
Pooled CRISPR screen design to identify modulators of cellular priming in the context of drug resistance to targeted therapies in melanoma. **A.** (top) In melanoma, the initial molecular profile of a cell (primed vs. un-primed) within an otherwise homogeneous population, indicated by green vs. gray coloring of cell, dictates the ultimate fate of the cell (e.g. proliferation vs. death) when exposed to therapy. Changing the number of cells in a given state (A, middle) can alter the number of resistant colonies that form upon addition of the BRAF^V600E^ inhibitor vemurafenib. **B.** We designed a pooled CRISPR screen to detect modulators of the cellular priming that leads to drug resistance. After transducing a library of single guide RNAs and expanding the population, we isolated cells with high expression of both NGFR and EGFR, then sequenced the single guide RNAs to determine which gene knockouts alter the frequency of these cells. Changes in the frequency of a given single guide RNA in this population (e.g. targets A and C) indicate that these targets may affect the frequency of NGFR^HIGH^/EGFR^HIGH^ cells in the population, and thus may affect frequency of cellular priming. **C.** After transducing a population of melanoma cells and isolating NGFR^HIGH^/EGFR^HIGH^ cells (see Fig. 1B), we quantified the frequency of each single guide RNA in the resulting population. Our screening scheme utilized three separate pooled single guide RNA libraries, one targeting epigenetic domains (top left), another targeting kinases (bottom left), and a final one targeting transcription factors (right). We organized the targets within each single guide RNA library by biological process. (While a given target could fall into several categories, each target is assigned to a single group and plotted only once.) Each dot represents a single guide RNA, grouped by gene target (5-6 single guide RNAs per target), with the log_2_ fold change representing the number of times the single guide RNA was detected in the sorted population versus an unsorted population of melanoma cells transduced with the same library. For display purposes, all single guide RNAs with fold changes beyond the axis limits were placed at the edge of the axis as indicated. For targets considered “hits” by our rubric (see methods), we labeled the single guide RNA dot by the color assigned to that biological process. Dots at the bottom of each pane correspond to non-targeting controls (single guide RNAs not targeting any loci in the genome) and cell viability controls (e.g. proteins required for cell survival and proliferation, but not specifically associated with rare-cell behavior). Supplemental Fig. 1 provides details on the effect of these editing controls.

With the advent of CRISPR/Cas9 technology, it is now possible to perform genetic screens to identify regulators of various molecular processes. For most cell fate transitions, including therapy resistance, virtually all screens have been designed to detect changes to the ultimate cellular fate only—i.e., changes in the final number of resistant cells, typically measured as a proliferation phenotype ^22–25^. However, an important aspect of plasticity is the process of stimulus-independent priming of rare cells in the population, which in this context is represented by the transient fluctuations in single cells that ultimately reprogram into a stably resistant cell fate ^14^. This priming processes may in principle have distinct regulatory mechanisms to that of the acquisition of resistance as a whole, presenting an opportunity to leverage screening techniques to specifically identify factors affecting cellular priming before the addition of drug. These factors may then also affect the overall degree of drug resistance, but potentially through new, previously undiscovered mechanisms that allow for new therapeutic targets that affect drug resistance in ways not revealed by classical resistance screens (Fig. 1A, bottom).

We here describe the results of genetic screens designed to capture modulators of single cell state variability that subsequently affect cell fate decisions. Specifically, in the context of melanoma, we performed pooled CRISPR/Cas9 genetic screens to discover modulators of the primed rare cell state that drives drug resistance. This new type of screen pointed to several new factors that affect the frequency of primed cells in melanoma populations, and ultimately, their resistance to targeted therapies. The transcriptome profiles induced by knocking out these factors revealed a novel mechanism that can increase or reduce drug resistance by increasing or decreasing the activity of differentiation pathways, respectively, as opposed to the more typical increased drug resistance induced by decreased differentiation. Drugs targeting these new mechanisms display a variety of synergistic effects when coupled with therapy, which can be dependent on the relative timing of drug application. Together, our results indicate that modulating cellular plasticity can alter cell fate decisions and may provide a new avenue for treating drug resistance.

## Results

### CRISPR/Cas9 genetic screens identify factors that affect primed cellular states

We wanted to identify factors that affected the fluctuations in cellular state that lead to single cells being primed to be drug resistant. We took advantage of a clonal melanoma cell line (WM989 A6-G3) that we have extensively characterized as exhibiting resistance behavior in cell culture that is broadly comparable to that displayed in patients ^14, 21, 26^. Phenomenologically, in cell culture, we observe that upon addition of a roughly cytostatic dose of the BRAF^V600E^ inhibitor vemurafenib (1µM), the vast majority of cells die or stop growing, but around 1 in 2,000-3,000 cells continues to proliferate, ultimately forming a resistant colony after 2-3 weeks in culture in vemurafenib. We have previously demonstrated that prior to the application of drug, there is a rare subpopulation of cells (pre-resistant cells) that express high levels of a number of markers, and that these “primed” cells are far more likely to become resistant than other cells ^14^. In order to identify modulators of the fluctuations that lead to the formation of this subpopulation of primed cells, we designed a large scale loss-of-function pooled CRISPR genetic screen (which we dubbed the “priming screen”) comprised of ∼13,000 single guide RNAs (sgRNAs) targeting functionally relevant domains of ∼2,000 proteins, with roughly six distinct single guide RNAs per domain (1402 transcription factor targets, 481 kinase targets, 176 epigenetic targets; each single guide RNA targets an important functional domain, see Supplemental Tables 1-3) ^27–29^.

To conduct the screen, we stably integrated *Streptococcus pyogenes* Cas9 (spCas9) into the WM989-A6-G3 cell line, creating the clonal line WM989-A6-G3-Cas9-5a3.

To screen for factors affecting cellular priming, we transduced this pooled library of single guide RNAs into this melanoma cell line. In order to ensure adequate sampling of the frequency of rare pre-resistant cells in the population, we expanded each cell in the culture to around 50,000-250,000 cells per each single guide RNA, resulting in a total screen size of roughly a billion cells. We then used a combination of magnetic sorting and flow cytometry to isolate cells that were positive for both EGFR and NGFR expression, both of which are markers of the primed cell subpopulation. We then sequenced the single guide RNAs in this sorted subpopulation to determine which single guide RNAs were over- or under-represented as compared to the unsorted total population. Here, over-representation suggests that knockout of the gene leads to an increased frequency of NGFR^HIGH^/EGFR^HIGH^ cells and vice versa (Fig. 1B). To select “hits” from the screen, we designed a series of criteria to identify and rank targets into confidence tiers (see methods for a detailed description of the selection criteria).

Our screen isolated several factors that affected priming for resistance. We obtained a set of 61 high confidence targets that affected the frequency of NGFR^HIGH^/EGFR^HIGH^ cells in our screen (Fig. 1C, Supplemental Table 4). Of these, 25 increased the frequency of NGFR^HIGH^/EGFR^HIGH^ cells, while the remaining 36 decreased the frequency. Beyond known factors in melanoma biology such as *SOX10* and *MITF* ^26, 30–32^, we identified several new factors not previously known to affect resistance to BRAF^V600E^ inhibition. These include *DOT1L*, which encodes an H3K79 methyltransferase associated with melanoma oncogenesis ^33^, and *BRD2*, which encodes a protein that is a member of the BET family, often overexpressed in human melanoma ^34^. We assessed the robustness and generality of our results through a secondary targeted screen in which 25 of the 34 high confidence targets tested replicated in the original WM989-A5-G3-Cas9 line. Furthermore, 20 of those 34 also replicated their effects in another melanoma line (451Lu-Cas9) (Supplemental Fig. 2, Supplemental Table 4). Together, these hits represented potential candidates for modulating therapy resistance by affecting cellular priming.

**Figure 2.**
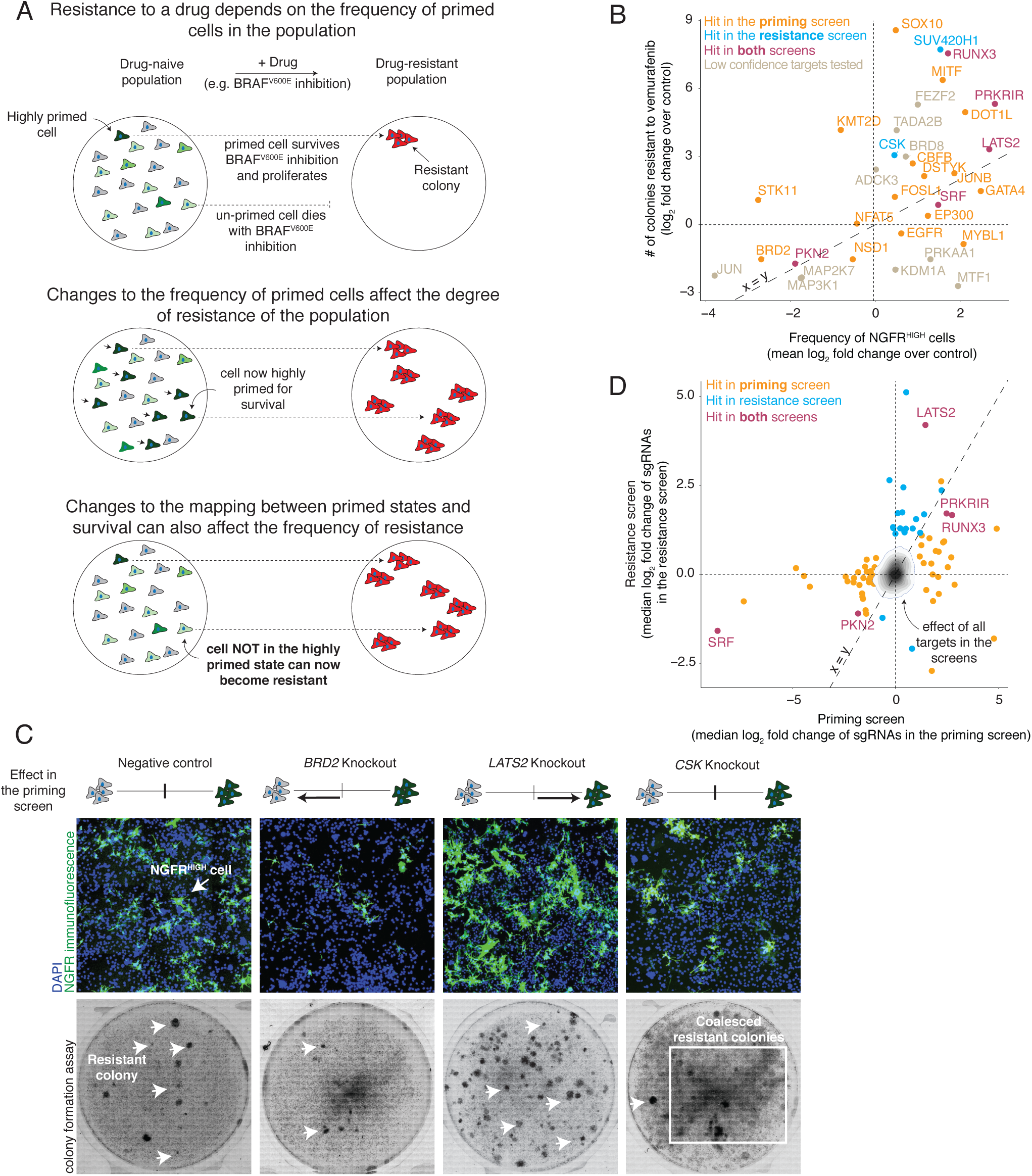
Effects of modulators of cellular priming on resistant colony formation. **A.** In melanoma, the frequency of primed cells in the population dictates the degree of resistance to BRAF^V600E^ inhibition. Changes to the mapping between cellular priming and a cell’s response to the drug can alter the number of resistant colonies that form upon addition of the BRAF^V600E^ inhibitor vemurafenib. **B.** Relationship between the frequency of NGFR^HIGH^ cells (x-axis) and the number of resistant colonies (y-axis). We plot the frequency of NGFR^HIGH^ cells as the mean log fold change over three replicates in the number of NGFR^HIGH^ cells following knockout of the gene indicated, normalized by cells with non-targeting sgRNAs. (For variability of the effect size across replicates of a given target, see Supplemental Fig. 5). We quantified the log_2_ fold change in the number of resistant colonies in the knockout cell line as compared to the non-targeting control cell lines. Orange points are targets identified as high confidence hits (Tier 1 and Tier 2) in the cellular priming screen; blue are those identified as high confidence hits in the resistance screen; purple are those identified as high confidence hits in both screens; gray, those that may have shown an effect in either or both screens, but were not classified as high confidence hits in either screen. **C.** To validate the phenotypic effect of targets identified by our genetic screens, we knocked out 83 of the targets and quantified both the frequency of NGFR^HIGH^ cells by immunofluorescence using anti-NGFR antibodies (top) and the number of resistant colonies (bottom) that form upon BRAF^V600E^ inhibition. Here we show example validation of BRD2 and LATS2 knockouts (hits in the cellular priming screen) and of CSK knockouts (hit in the resistance screen only). The schematic represents the effect of the knockout in the priming screen on the frequency of NGFR^HIGH^/EGFR^HIGH^ cells. **D.** Effect overlap between hits from the cellular priming and resistance screens. Each target’s position (dots) represents the number of times (as median log2 fold change) the single guide RNAs were detected in the NGFR^HIGH^/EGFR^HIGH^ population vs. an unsorted population of melanoma cells (priming screen, x-axis), or in the population of cells resistant to vemurafenib vs. the population of cells prior to treatment (resistance screen, y-axis). Orange labels correspond to high confidence targets (Tier 1 and Tier 2) in the cellular priming screen; blue corresponds to high confidence targets in the resistance screen; purple corresponds to high confidence targets in both screens. The effects of all targets in both screens are displayed as a density histogram.

### Changes in drug resistance can occur by priming-dependent and independent mechanisms

Our priming screen was designed to isolate candidates that affected the frequency of cells that were in the primed cellular state *before* the addition of drug. Conceptually, it is also possible that there may be a distinct set of factors that can affect the overall rate of resistance *without* affecting the frequency of primed cells. It is useful here to separate the notion of priming, which we use to refer to the cellular *state* associated with high levels of expression of resistance (i.e., NGFR^HIGH^ cells), from the notion of pre-resistance, which is the set of cellular states that, upon adding drug, will eventually develop into a stably resistant colony. The former definition is dependent on the state of the cell, whereas the latter is dependent on the drug: for instance, if one added a lower concentration of drug that allowed for more resistant colonies to form, the initial primed states would remain unchanged, but the number of pre-resistant cells would change because more cells became resistant. In this framework, then, one could imagine that any particular factor could increase resistance either by increasing the frequency of primed cells (Fig. 2B, middle) or by allowing more cells that are partially primed to continue to grow upon addition of drug and become resistant; i.e., lowering the putative threshold (Fig. 2A, bottom) (or both).

We wanted to measure the degree to which the factors we identified could affect both priming and resistance via these two different mechanisms. First, because our first screen was designed to identify factors affecting priming, we also ran another genetic screen (as well as a secondary targeted screen with another melanoma cell line) with the ultimate readout being number of resistant colonies; i.e., a conventional survival screen (“resistance screen”) (Supplemental Figs. 3-4, Supplemental Table 4). We identified 20 high confidence factors that, when knocked out, increased the number of resistant cells and 4 that reduced the number of resistant cells. These included factors involved in signaling pathways like MAPK (*CSK*) ^35^, Wnt/B-catenin (*KDM2A*) ^36^, and Hippo (*LATS2*) ^37^.

**Figure 3.**
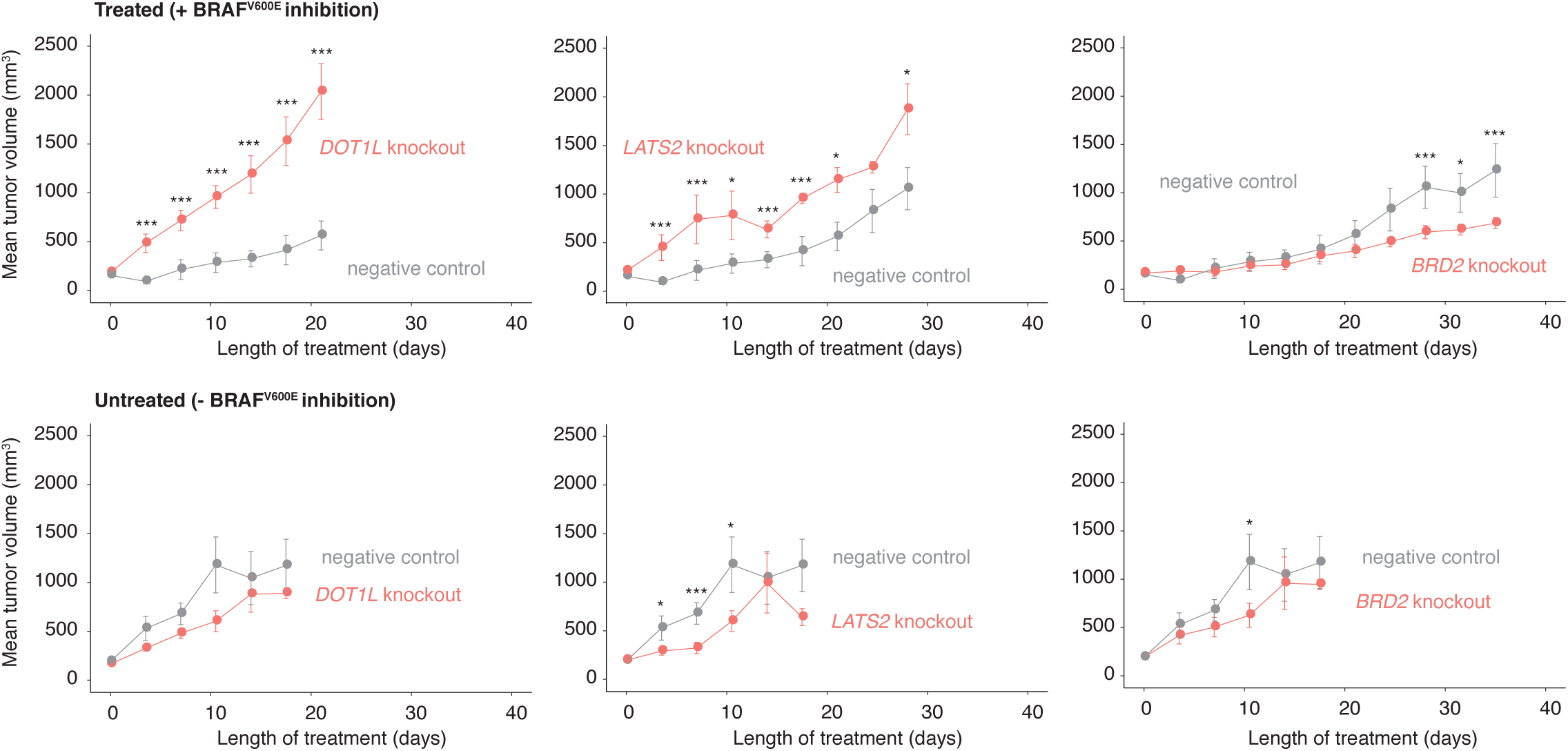
Effect of modulators of cellular priming on growth of BRAF^V600E^-resistant tumors in vivo. A. Tumor volume as a function of time in patient-derived xenografts (NOD/SCID mice) treated with a BRAFV600E inhibitor (top) or vehicle control (bottom). Here, we inject each mouse with DOT1L-, LATS2-, or BRD2-knockout WM989-A6-G3-Cas9 cells (orange) or with the same cell line without a gene knockout (gray). The values plotted represent the mean tumor volume across mice caryying the same knockout. Error bars represent the standard error of the mean. *** are timepoints at which the difference in tumor volume between knockout and control groups reached p ≤ 0.05. Similarly, * represents p = 1 ≤≥ 0.05 (see methods). Each group started with n = 6 mice, and we plotted the mean tumor volume up until both knockout and negative control groups have at least n = 3 each.

**Figure 4.**
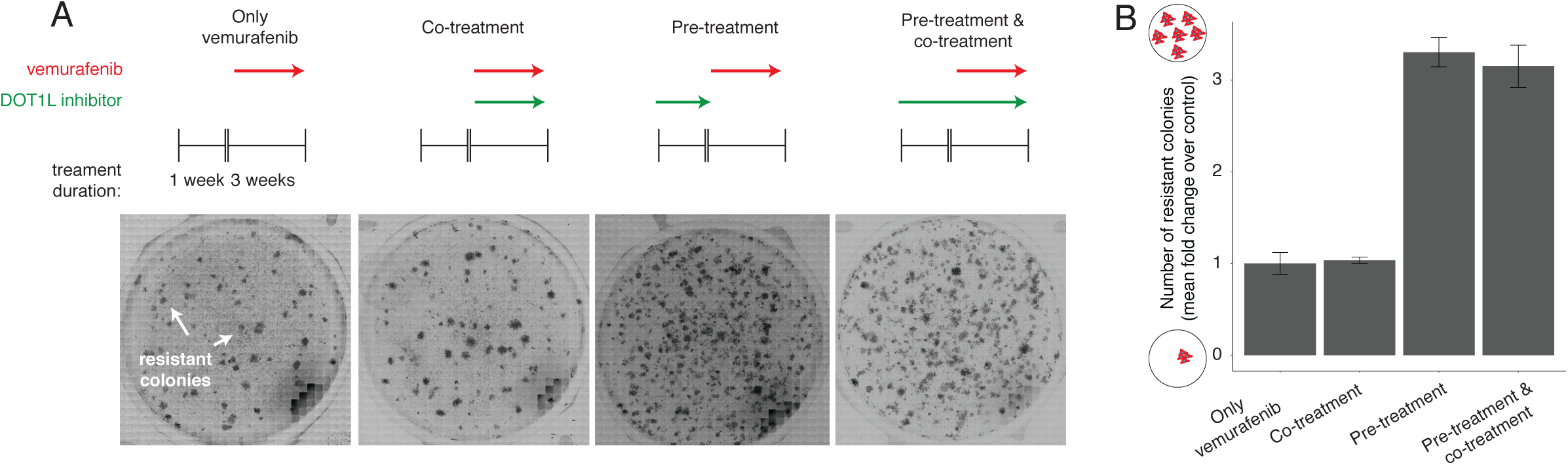
Effect of targeting cellular priming at different times relative to BRAF^V600E^ inhibition. **A.** To assess the effect of DOT1L inhibition (green arrows, pinometostat at 4µM) at different times on a cell’s ability to survive BRAF^V600E^ inhibition, we first established a baseline number of colonies that grow when WM989-A6-G3 cells are exposed to 1µM of vemurafenib for three weeks (leftmost panel). Then, in a separate population, we either inhibited BRAF^V600E^ and DOT1L simultaneously (co-treatment), inhibited DOT1L first (seven days) and then BRAF^V600E^ (three additional weeks; pre-treatment), or inhibited DOT1L before (seven days) and during three weeks of vemurafenib treatment (pre-treatment and co-treatment). **B.** Number of resistant colonies that result from each therapeutic regimen in Fig. 2C as the mean fold change over baseline (vemurafenib alone) for three replicates normalized to the number of cells in culture prior to BRAF^V600E^ inhibition. Error bars indicate the standard error of the mean over triplicates.

The priming and resistance screens were designed to probe distinct biological behavior, and so we predicted that of the factors identified, some would affect the frequency of priming and some would affect the putative “threshold” that must be surpassed for the acquisition of resistance (and some may affect both). To systematically evaluate whether such differences existed, we directly looked on a knockout-by-knockout basis for changes in the frequency of primed cells (by NGFR immunofluorescence) in 83 different targets from both the priming and the resistance screens, and further looked for changes in resistance (by measuring the number of resistant colonies produced) in 35 of those (Fig. 2B, Supp. Fig. 5). We found that these individual knockouts exhibited a range of changes in both the frequency of NGFR^HIGH^ cells and the number of resistant colonies formed. Firstly, many hits from the priming screen (15/21 tested by both immunofluorescence and resistant colony formation) showed the predicted increase and decrease in both frequency of priming and concomitant changes to the number of resistant colonies (e.g. LATS2, BRD2, respectively; Fig. 2C). This result demonstrates that hits that led to changes in the frequency of cells expressing *NGFR* were associated with changes in priming and, consequently, resistance.

**Figure 5.**
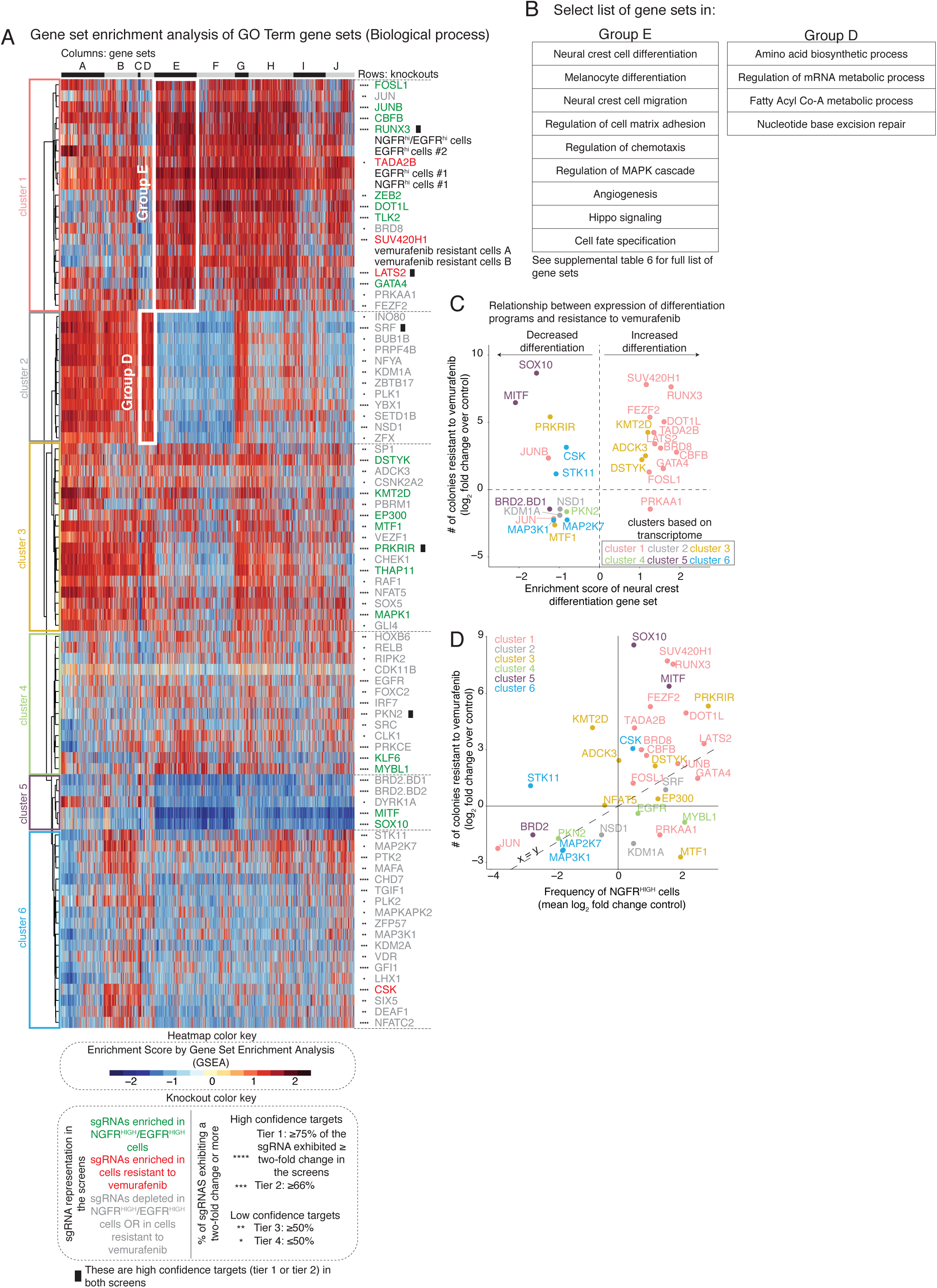
Gene set enrichment analysis of the transcriptional effects induced by the knockout of select screen targets. **A.** The heatmap represents biclustering analysis of different knockout cell lines (rows) based on the Gene Set Enrichment Analysis score of Gene Ontology gene sets (Biological process GO terms, columns in heatmap). Within the heatmap, red indicates enrichment in the sense that there are more differentially upregulated genes in knockout vs. control in that gene set than expected by chance, whereas blue indicates enrichment of downregulated genes (shade indicates degree of enrichment). Each target knockout (rows) represents transcriptomes of biological triplicates (unless otherwise stated on Supplemental Table 4). Target labels (rows) in green indicate genes whose knockout increased the frequency of NGFR^HIGH^/EGFR^HIGH^ cells in the screen, while red indicates targets whose knockout increased the number of cells resistant to vemurafenib, and gray indicates targets that decreased the frequency of either NGFR^HIGH^/EGFR^HIGH^ cells or of cells resistant to vemurafenib. As before, we organized targets into high confidence hits (Tier 1 and Tier 2) and low confidence hits (Tiers 3 and Tier 4) based on the percentage of single guide RNAs against a target that showed at least a two-fold change in the initial screen (see knockout color key).The asterisks next to the label indicate the tier (Tier 1, ****; Tier 2, ***; Tier 3, **; Tier 4, *). Information regarding validation rates of each tier can be found in the supplemental figures 8 and 9. Based on the dendrogram on the left, we grouped targets into six clusters. We also clustered gene sets (columns) into groups, labeled by the letters on top of the heatmap. The white boxes inside the heatmap demark groups of gene sets specifically upregulated in a given cluster. **B.** Select list of gene sets in groups D and E from Figure 5A. For a complete list of gene sets within each group, see supplementary table 6. **C.** Relationship between the expression of genes involved in neural crest differentiation (x-axis) and the number of colonies resistant to vemurafenib (y-axis) following the knockout of a target. For each knockout, we plot the expression of neural crest differentiation genes as the enrichment score obtained through gene set enrichment analysis for the neural crest differentiation gene set (GO term). We quantified the log_2_(fold change) in the number of resistant colonies in the knockout cell line as compared to the non-targeting control cell lines. Colors represent the cluster grouping of each knockout based on Figure 5A. **D.** Relationship between the frequency of NGFR^HIGH^ cells (x-axis) and the number of resistant colonies (y-axis). We plot the frequency of NGFR^HIGH^ cells as the median log (fold change) over three replicates in the number of NGFR^HIGH^ cells following knockout of the indicated gene normalized by cells with non-targeting single guide RNAs. (For variability of the effect size across replicates of a given target, see Supplemental Fig. 5.) We quantified the log_2_(fold change) in the number of resistant colonies in the knockout cell line as compared to the non-targeting control cell lines. We color-coded all targets by groupings based on their transcriptomes (see Fig. 5A) following knockout of the gene indicated.

However, while the general trend indicated such a pattern, knockouts of many genes varied widely in the degree to which this relationship held (Fig. 2B). For instance, knockout of *EP300* resulted in a ∼two-fold increase in the number of NGFR^HIGH^ cells but only a small increase in the number of resistant colonies, while knockout of *CSK* resulted in only a small increase in the number of NGFR^HIGH^ but had at least a six-fold increase in the number of resistant colonies (Fig. 2B-C). (Importantly, the number of colonies for the CSK knockout is an underestimate due to difficulties in accurately counting colonies in highly-confluent plates, see Supplemental Fig. 5C for zoomed image of colonies, and is most likely why *CSK* was a dominant hit in our resistance screen.) Conceptually, hits like *CSK* could operate by lowering the putative threshold of priming that cells need to surpass in order to progress to acquire resistance *without* changing the degree of priming in individual cells, thus leading to more resistant colonies without affecting priming. Another possibility is that knocking out *CSK* does change priming, but in a way that is not reflected in our measurements, that is, in a change in NGFR expression. While our results cannot conclusive exclude this latter possibility, some of our results argue in favor of changes to the threshold. For instance, the *CSK* knockout cell line showed an increase in the number of resistant colonies but also an increase in the number of resistant cells that do not form colonies (Fig. 2C, Supplemental Fig. 5C). This suggested that, in addition to the usual pre-resistant cells that form colonies, an additional set of cells in the *CSK* knockout line were now enabled to survive drug. This suggests that the “threshold” for cells to survive drug may have changed; i.e., the mapping between the degree of priming and the ultimate resistant fate has been altered by the removal of *CSK*.

Notably, of the factors identified by the resistance screen, only 5 were also identified in our priming screen (Fig. 2D). In principle, if it were possible to run the resistance screen to saturation—i.e., isolate *all possible* factors affecting resistance—then the resistance screen should be able to find all priming factors that affect resistance. However, in practice, the number of cells required make it very difficult to run these screens to true saturation, and thus it is possible that dominant hits that change resistance alone (e.g. *CSK*) comprised so many cells in the pooled resistance screen that other hits associated with changes in priming became difficult to detect. This possibility highlights the potential of screens targeting priming to reveal novel categories of hits that may otherwise elude detection.

### Changes in the frequency of primed cellular states lead to differences in tumor growth *in vivo*

We found that the factors we identified that modulate cellular priming can further lead to differences in overall resistance to BRAF^V600E^ inhibition in cell lines, but we still wondered whether these same factors can affect resistance in an *in vivo* setting, in which complex microenvironmental factors may also affect therapy resistance ^38^. We thus tested whether knocking out three of the factors isolated from our cellular priming screen affected resistance *in vivo*: *DOT1L* and *LATS2*, which increased the frequency of NGFR^HIGH^/EGFR^HIGH^ cells *in vitro*, and *BRD2*, which decreased the frequency of NGFR^HIGH^/EGFR^HIGH^ cells *in vitro* (all three also exhibited corresponding differences in resistance to BRAF^V600E^ inhibition *in vitro*).

After knocking out these targets in WM989-A6-G3-Cas9-5a3, we injected the cells into NOD/SCID mice (n=12 mice per knockout) and allowed tumors to develop. We quantified the tumor volume in each mouse over time, comparing tumors that developed in mice injected with similar cells (WM989-A6-G3-Cas9-5a3) but without any gene knocked out (Fig. 3). Overall, we observed patterns consistent with our *in vitro* results: at the treatment endpoint (see methods) *DOT1L* knockout tumors treated with a BRAF^V600E^ inhibitor produced tumors that were roughly 3.5 times larger than controls (p = 0.010), and *LATS2* knockout cell lines produced tumors that were 1.6 times larger than controls (p = 0.062). On the other hand, the mice that received melanoma cells with the *BRD2* knockout had tumors that were approximately half as big as controls (p = 0.045). (In the absence of drug, both knockout and control melanoma cells showed roughly similar growth dynamics (Fig. 3, bottom)). Overall, our results demonstrate that the factors isolated by our cellular priming screen also affect the response of tumors to BRAF^V600E^ inhibition *in vivo*.

### Relative timing of targeting variability can affect drug resistance

The fact that many of the factors we identified had different effects on priming vs. full acquisition of resistance as measured by resistant colony formation suggested that these factors may work by different mechanisms, and that these mechanisms may potentially interact or override each other in complex ways dependent on relative timing. For instance, a factor that affects specifically priming could affect the number of cells in the pre-resistant state, but once cells are subjected to before BRAF^V600E^ inhibition and begin reprogramming towards stable resistance, the factor may no longer have any effect. In such a case, inhibiting this factor *before* the adding the BRAF^V600E^ inhibitor would be critical.

To test for such a possibility, we used the DOT1L inhibitor pimenostat ^39, 40^ (which increases the number of colonies resistant to vemurafenib over a range of doses; Supplemental Fig. 7A) to see if timing of DOT1L inhibition would affect the formation of resistant colonies. In addition to the standard vemurafenib treatment, we both pre-treated with the DOT1L inhibitor for seven days before adding vemurafenib and co-treated with the DOT1L inhibitor concurrently with vemurafenib (we tested both pre-treatment followed by vemurafenib alone and pre-treatment followed by concurrent treatment) (Fig. 4A). We found that pre-inhibition of DOT1L resulted in three-fold more colonies than with BRAF^V600E^ inhibition alone, but that co-treatment with the DOT1L and BRAF^V600E^ inhibitors led to no change in the number of resistant colonies (Fig. 4B), suggesting that DOT1L inhibition is altering the distribution of states of the cells, and consequently the number of cells that develop resistance to BRAF^V600E^ inhibition. Our results demonstrate that the relative timing of inhibition of cellular priming vis a vis mainline therapy can have a profound effect on resistance.

### Knockout of novel genes that increase the frequency of primed cell states also increase cellular differentiation

Our screens revealed a large number of factors affecting priming that act across a range of biological processes, including a variety of signaling pathways and transcriptional regulatory mechanisms. Interestingly, *a priori*, no particular pathway appeared to dominate the set of identified factors; however it is possible that seemingly unrelated genes nevertheless affect priming through common biological processes.

To look for such commonalities, we used RNA sequencing to measure genome-wide transcript abundance levels for 266 knockout cell lines targeting 80 different proteins taken from both the priming and resistance screens (each targeted with 2-3 separate single guide RNAs; see supplementary table 4), reasoning that if two genes participated in a particular biological process, then the transcriptomes associated with knocking them out may exhibit similar patterns of differential expression.

Initially, we clustered the transcriptome profiles from the different cell lines, including only genes differentially expressed in at least one sample (Supplemental Fig. 10A). We found that while the transcriptomes induced by some gene knockouts were clearly distinct (such as *MITF*, *SOX10* and *KDM1A*), many others appeared to show only relatively small differences from the parental cell line, despite the fact that our validation results showed that these knockouts exhibited clear effects on the resistance potential of the population. We thus reasoned that while the sets of genes whose expression change in our knockouts may be non-overlapping, these genes could still belong to similar categories of biological processes; i.e., different knockouts may all affect different genes all within a common pathway, for instance differentiation. Thus, using the transcriptome of each knockout, we performed a gene set enrichment analysis (GSEA, see methods) and obtained an enrichment score for a number of biological processes from the Gene Ontology terms database (Fig. 5A) ^41^. Using these enrichment scores, the knockout lines clustered in a more obvious pattern. Notable clusters include cluster 5, containing the canonical melanocyte master regulators *MITF* and *SOX10*, and cluster 1, containing *DOT1L*, *LATS2*, *RUNX3* and *GATA4*.

Interestingly, knocking out MITF and SOX10 increases drug resistance, as does knocking out most members of cluster 1, but the transcriptome profiles of these two clusters appeared to be roughly opposite of each other. We inspected the GO gene sets in Group E, which appeared maximally different between MITF/SOX10 and cluster 5, and found that these gene sets included several related to differentiation, including sets for melanocyte differentiation and neural crest differentiation (Fig. 5B). The knockout of MITF and SOX10 appeared to decrease the expression of these genes, matching the general consensus that drug resistance is typically driven by dedifferentiation ^8, 26^. In that context, the finding that most elements of cluster 1 increased resistance by further promoting differentiation was unexpected (Fig. 5C), suggesting a possible novel mechanism by which one could affect drug resistance; the latter has further support from our findings using the DOT1L inhibitor (Fig. 4). This axis of differentiation was coordinated across several gene sets, as revealed by principal components analysis of the expression heat map (Supplemental Fig. 10B). (Note that the role of *MITF* in therapy resistance is complex in general ^42^).

Clusters of targets that lead to different degrees of differentiation also seem to correspond to distinct phenotypic profiles, meaning the resultant changes in the frequency of NGFR^HIGH^ cells and number of resistant colonies. For instance, the transcriptomes of the knockouts in cluster 1 seem to mimic many aspects of the transcriptomes of NGFR^HIGH^, EGFR^HIGH^, NGFR^HIGH^/EGFR^HIGH^, and even vemurafenib resistant melanoma cells (e.g. high expression of genes involved in cell-matrix adhesion, angiogenesis, and cell migration; Fig. 5A,B). Knockout of these targets showed a strong correspondence between the frequency of NGFR^HIGH^ cells and the number of colonies that developed under BRAF inhibition, suggesting that the increase/decrease in the frequency of pre-resistant cells was the cause of increased/decreased resistance (Fig. 5D). Often, this relationship was relatively proportional, as was the case for the knockout of *LATS2, JUNB, FOSL1,* and *CBFB*. For *MITF* and *SOX10* (cluster 5), however, the relationship between the frequency of NGFR^HIGH^ cells and the number of resistant colonies was much weaker, with very large changes in the latter but not the former. Accordingly, our transcriptomic analysis suggests that these knockouts lead to changes in gene expression that are distinct from those of NGFR^HIGH^/EGFR^HIGH^ cells.

The transcriptome analysis also revealed different categories of knockouts that resulted in a *reduction* (as opposed to increase) of the number of resistant colonies. Some resistance reducing knockouts (*BRD8* and *PRKAA1*) clustered with *DOT1L*, while another (*BRD2)* clustered with *MITF/SOX10*. It is possible that these factors work in inverse ways to reduce drug resistance by either affecting differentiation or dedifferentiation. Meanwhile, the majority of resistance reducing knockouts appeared to cluster separately into distinct clusters, generally through changes in the expression of a distinct set of genes. For one cluster (cluster 2), the set of genes whose expression was affected included several associated with metabolism (e.g. biosynthesis of amino acids and Acyl Co-A metabolism), suggesting that modulation of metabolic processes may be a means of reducing drug resistance (Supplemental Table 6). The other clusters did not show any coherent set of biological processes affected (e.g. *SRC*, *IRF7*, *PKN2*, among others), rendering that particular pathway or set of pathways rather mysterious.

## Discussion

We have here demonstrated, using high-throughput genetic screening, that there are genetic factors that can alter cellular plasticity in cancer cells, thereby affecting their resistance to targeted therapeutics. We identified a variety of new factors that appear to work through new pathways that can affect therapy resistance in novel, time-dependent ways. These factors revealed new possible vulnerabilities that a conventional genetic screen targeting resistance did not uncover, thus demonstrating the potential for screens specifically designed to target single cell variability to reveal new biological mechanisms that may subsequently emerge as therapeutic opportunities. Drug screens targeting gene expression “noise” have also shown similar therapeutic potential ^43^.

While we isolated several new factors that specifically affected cellular variability, it is important to note that no single factor we isolated resulted in a change in cellular variability that was stronger than all the rest; i.e., no factor emerged as the “smoking gun”. This may be the result of the fact that our screen did not target all potential regulators. Alternatively, it may be that the biology of cellular variability is intrinsically multifactorial, with the coherent activity of many factors being required for cells to ultimately enter the highly deviated cellular state responsible for phenotypes like drug resistance ^15^. Larger scale screens may help reveal a more complete picture of the origins of rare cell behavior; however, the limitations imposed by the rarity of the pre-resistant cellular phenotype make this rather difficult. The raw numbers of cells required to properly sample these rare cell behaviors in a pooled genetic screening format remains a major technical challenge for the field of rare cell biology.

Indeed, it is the very difficulty of performing these screens at full depth that provides motivation for screening for variability rather than simply screening for resistance. If one is primarily interested in factors affecting resistance, then in principle such a screen, if carried to saturation, would reveal all such factors, including those that exert such an effect via modulation of cellular variability. However, the degree of overlap in the factors identified between our variability screen and our conventional resistance screen was relatively small. This lack of overlap suggests that distinct biological processes may dominate the results of these differently designed screens. That of course in turn raises the question of why one might want to perform variability screens at all, given that the phenotype of interest is resistance. Our results on timing of variability inhibition suggest that while the mechanisms governing rare cell variability may not appear as potent as those revealed by conventional resistance screens, the fact that they represent *distinct* mechanisms means that they may present an opportunity to be used in tandem. It is also possible that these mechanisms may be more dominant in other, more clinically relevant contexts.

In our validation studies, for several factors, we measured the effects of knocking out those factors on both the number of NGFR^HIGH^ cells (which serves as a proxy for the primed cellular state) and the number of resistant colonies upon adding vemurafenib. Interestingly, different knockouts affected both of these validation metrics differently, with some (e.g. *LATS2*) both increasing the frequency of NGFR^HIGH^ cells as well as concomitantly increasing the number of resistant cells, and some (e.g. *CSK*) dramatically increasing the frequency of resistant cells without a proportional change in the frequency of NGFR^HIGH^ cells. One possible way to conceptualize these distinct phenotypic outcomes is that the former category of knockout affects primarily cellular priming, i.e., the cellular state, while the latter affects the mapping between these primed states and their fates upon addition of vemurafenib. In one simple model, one could imagine a distribution of cellular states in the initial population and a threshold whereby cells above the threshold survive the drug and those below the threshold do not (Fig. 6). In this model, some knockouts may alter the distribution of cells in the initial population, thus rendering a different proportion of them above or below the threshold, or may alter the threshold itself, or potentially some combination of both. It is wise to caution against this simple interpretation, however. First, we note that NGFR expression is just a marker for the pre-resistant state, and it may be that factors may affect the frequency of pre-resistant cells without showing any effect on NGFR expression, thus giving the false appearance of a change in the mapping. (Arguing against this, however, is the fact that the transcriptomes of knockouts such as *DOT1L* that increase the frequency of NGFR and resistance appear to be similar to the profile of NGFR^HIGH^ cells themselves; Fig. 5A). Further molecular profiling of individual cells from these knockouts may help reveal the ways in which the molecular state of these cells changes. Secondly, it is also likely that the categorization of fates as “resistant” or “dead” is dramatically oversimplified, and that there may be a number of different types of resistant cells (anecdotally, we have noticed that the resistant cells in some of our knockout lines do appear morphologically different from those formed in the unperturbed cell line). Such results suggest that there is a mapping from a continuum of initial cellular states to multiple, canalized, or even potentially continuous cellular fates. An important future direction is to characterize this mapping and its regulation.

**Figure 6.**
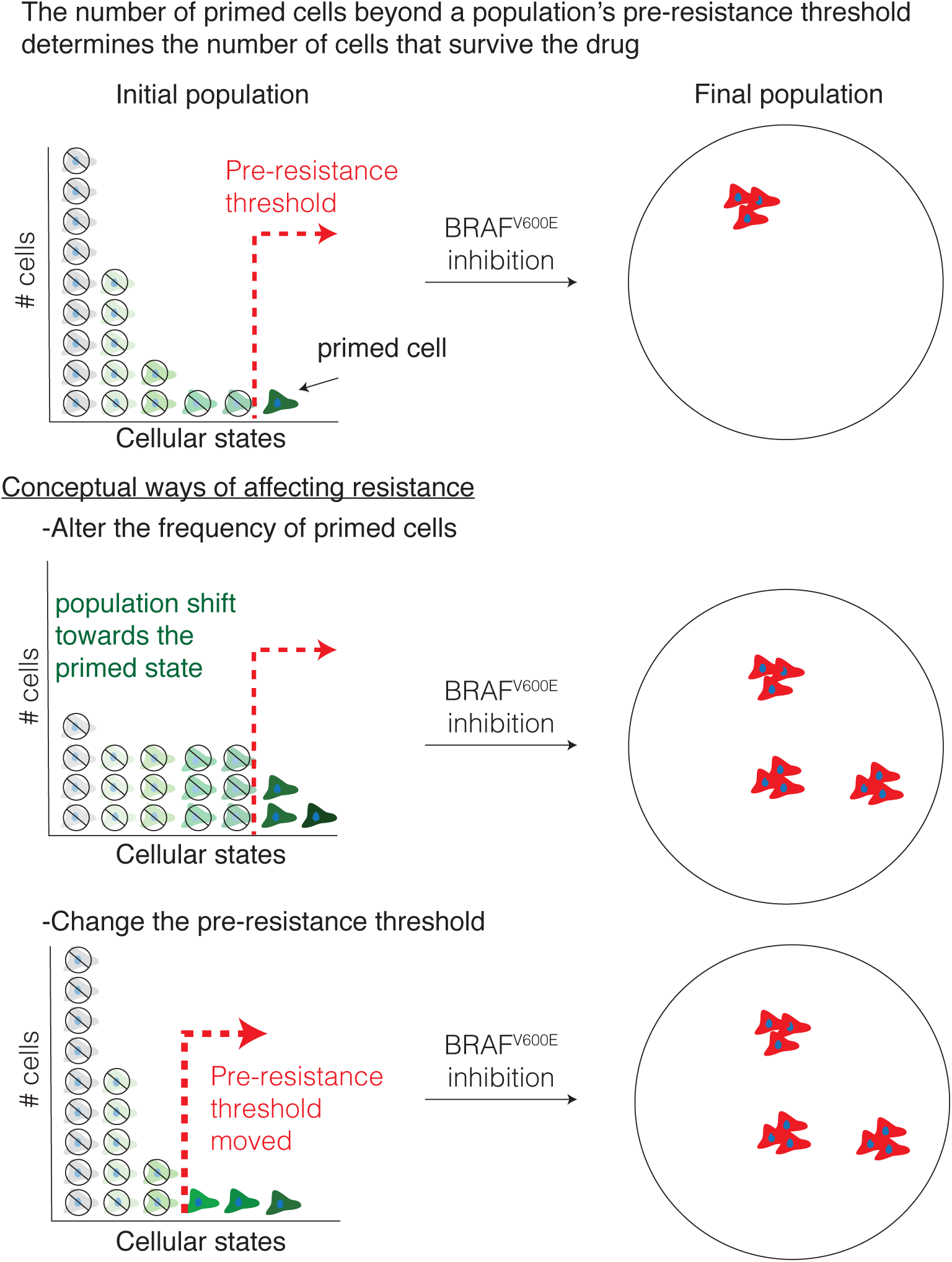
Model of pre-resistant threshold and cellular priming in the development of resistance to targeted therapies. Variability in the expression of various markers are associated with an individual cell’s probability to survive drug treatment. In one simple model, cellular variability occurs along a single ordinate, which can be conceptualized as the degree of “greenness”. In this model, there is a threshold (red line, top panel) that divides cells along this axis into those that adapt to the drug and become resistant vs. those that no longer proliferate when challenged with drug. Here, there are at least two ways by which one could conceivably alter the number of cells that survive the drug. In one scenario (middle) the distribution of “greenness” could change, leading to more cells being above the threshold, leading to more resistant colonies. In another scenario, the distribution of phenotypes remains unchanged, but the threshold itself moves, also leading to more resistant colonies. Our results suggest (but do not prove) that both scenarios may play out to varying degrees as a result of different genes being knocked out.

Here, we have focused on cellular variability in the context of drug resistance in cancer. However, we have observed similar rare-cell variability in primary melanocytes ^14^, raising the possibility that the same variability may play a role in normal biological processes as well. It is thus possible that the factors we have isolated may play a role in regulating variability in these normal biological contexts, and it remains to be seen whether such factors act primarily in melanocytes or act more generally across different cell types in various tissues. Indeed, we believe variability will emerge as a key aspect of cellular plasticity in general, and that framing plasticity as a mapping between variable cellular states and ultimate phenotypic fates may prove a fruitful conceptual framework.

## Acknowledgements

We want to thank Dr. Meenhard Herlyn for always providing excellent advice and guidance. We also thank the Flow Cytometry core, especially Florin Tuluc, at CHOP for all their advice and help. We also thank all members of the Raj Lab as well as John Murray for their comments and suggestions. We thank C. Vakoc for providing the transcription factor, epigenetic regulator, and kinase domain-focused sgRNA library. J.S acknowledges support from Linda Pechenik Montague Investigator Award and Cold Spring Harbor Laboratory sponsored research. AR acknowledges NIH/NCI PSOC award number U54 CA193417, NSF CAREER 1350601, P30 CA016520, SPORE P50 CA174523, NIH U01 CA227550, NIH 4DN U01 HL129998, NIH Center for Photogenomics (RM1 HG007743), and the Tara Miller Foundation. AW acknowledges support from CA207935 and CA174746. AW acknowledge support from CCSG P30CA010815 and NIH U01 CA227550.

## Author contributions

E.T., J.S., and A.R. designed and supervised the study. E.T. performed the experiments and analysis. E.A., K.B. assisted with CRISPR screens. S.B. assisted with tissue culture, image acquisition, and analysis. L.B. designed image analysis software. M.F., G.A. performed in vivo assay. B.E, S.S. assisted with acquisition of transcriptomic data. B.E, S.S., I.M. assisted with data analysis. A.W. provided guidance on interpretation of the data.

## Supplemental Materials

### Material and Methods

#### Cell Culture

We obtained patient-derived melanoma cells (WM989 and 451Lu, female and male, respectively) from the lab of Meenhard Herlyn. For WM989 we derived a single cell subclone (A6-G3) in our lab ^14^. We grew these cells at 37°C in Tu2% media (78% MCDB, 20% Leibovitz’s L-15 media, 2% FBS, and 1.68mM CaCl2). We authenticated all cell lines via Human STR profiling. We periodically tested all cell lines for mycoplasma infections.

#### Plasmid Construction and single guide RNA Cloning

All the Cas9 positive melanoma cell lines in this study were derived by lentiviral transduction with a Cas9 expression vector (EFS-Cas9-P2A-Puro, Addgene: 108100). All the single guide RNAs were cloned into a lentiviral expression vector LRG2.1(Addgene: #108098), which contains an optimized single guide RNA backbone. The annealed single guide RNA oligos were T4 ligated to the BsmB1-digested LRG2.1 vector. To improve U6 promoter transcription efficiency, an additional 5’ G nucleotide was added to all single guide RNA oligo designs that did not already start with a 5’ G.

#### Construction of Domain-Focused single guide RNA Pooled Library

Gene lists of transcription factors (TF), kinases, and epigenetic regulators in the human genome were manually curated based on the presence of DNA binding domain(s), kinase domains, and epigenetic enzymatic/reader domains. The protein domain sequence information was retrieved from NCBI Conserved Domains Database. Approximately 6 independent single guide RNAs were designed against individual DNA binding domains (Supplementary tables 1-3).^27–29^ The design principle of single guide RNA was based on previous reports and the single guide RNAs with the predicted high off-target effect were excluded (Hsu et al. 2013). For the initial pooled CRISPR screens, all of the single guide RNAs oligos including positive and negative control single guide RNAs were synthesized in a pooled format (Twist Bioscience) and then amplified by PCR. PCR amplified products were cloned into BsmB1-digested LRG2.1 vector using Gibson Assembly kit (NEB#E2611). For the targeted pooled validation screens, individual single guide RNAs were synthesized, cloned, and verified via Sanger sequencing in a 96-well array platform (Supplementary table 5). Individual single guide RNAs were pooled together in an equal molar ratio. To verify the identity and relative representation of single guide RNAs in the pooled plasmids, a deep-sequencing analysis was performed on a MiSeq instrument (Illumina) and confirmed that 100% of the designed single guide RNAs were cloned in the LRG2.1 vector and the abundance of >95% of individual single guide RNA constructs was within 5-fold of the mean (data not shown).

#### Lentivirus preparation

We produced lentivirus containing single guide RNAs using HEK293T cells cultured in DMEM supplemented with 10% Fetal Bovine Serum and 1% penicillin/streptomycin. When the cells reached 90-100% confluency, we mixed the single guide RNA vectors with the packaging vector psPAX2 and envelope vector pVSV-G in a 4:3:2 ratio in OPTI-MEM (ThermoFisher Scientific: #31985070) and polyethylenimine (PEI, Polysciences: #23966). We collected viral supernatants for up to 72 hours twice daily.

#### Transduction of spCas9

We introduced the stable expression of spCas9 via spinfection of lentivirus along with 5ug/ml polybrene for 25 minutes at 1750 rpm. We exchanged the media ∼6 hours post-transduction and selected for cells expressing spCas9 via puromycin selection (1-2μg/ml, 1 week). For WM989-A6-G3, we generated two cell lines, WM989-A6-G3-Cas9 and WM989-A6-G3-Cas9-5a3, the later being a single cell isolate of the bulk Cas9-expressing population. We verified that this cell line was capable of editing the genome and that it still contained pre-resistant cells marked by the expression of drug-resistance markers (Supplemental Fig. 11). Following the same methodology, we generated a 451Lu-Cas9 cell line from 451Lu cells.

#### Transduction of lentivirus containing single guide RNAs

For transfection of melanoma cells, we infected cells with lentivirus and 5ug/ml polybrene for 25 minutes at 1750 rpm. We exchanged the media ∼6 hours post-transfection. We quantified the percent of the population transfected by measuring the number of GFP-positive cells at day 5 post-transfection. For the screens, we aimed to transfect 30% of the population. For all other experiments, we aimed to transfect >95% of the population.

#### Initial pooled CRISPR screens

We worked with three main pooled single guide RNA libraries in WM989-A6-G3-Cas9-5a3 cells. These libraries targeted ∼2,000 different kinases, transcription factors, and proteins involved in epigenetic regulation. In total, the libraries contained ∼13,000 different single guide RNAs including non-targeting and cell-viability editing controls (Supplementary tables 1-3). We aimed to transfect > 1,000 cells per single guide RNA and isolated ∼1,000 cells per single guide RNA about a week post-transfection and prior to any selection. These baselines allowed us to validate the efficiency of our screen by single guide RNA enrichment/depletion of non-targeting controls and of controls that affect cell viability (Supplemental Fig. 1). Additionally, these baselines helped us identify single guide RNAs with lethal effects in our cells. Given that we were interested in rare cell phenotypes that exist in 1:2000 cells or less, throughout our screens we significantly expanded the population of cells to 50,000-250,000 cells per single guide RNA, often surpassing a billion cells per screen. This scale allowed us to observe the rare cell phenotypes dozens-to-hundreds of times in each of our controls (and in each of our single guide RNAs).

The priming screen aimed to identify perturbations that altered the frequency of NGFR^HIGH^/EGFR^HIGH^ cells. To this end, one month after we transfected and expanded the cells, we isolated the NGFR^HIGH^/EGFR^HIGH^ cells via magnetic cell sorting (MACS) followed by fluorescence-activated cell sorting (FACS) (see below). We also collected an additional ∼1,000 cells per single guide RNA, without any selection, for comparison. Then, we isolated DNA from the cells and built sequencing libraries (see below) to quantify the representation of each single guide RNA in the NGFR^HIGH^/EGFR^HIGH^ population and compare it to the unsorted baseline.

In the resistance screen we aimed to identify proteins important for the development of resistance to vemurafenib. Here, we treated the cells as above, except that instead of isolating NGFR^HIGH^/EGFR^HIGH^ cells we grew cells resistant to vemurafenib (see below) by exposing the cells to vemurafenib for three weeks. As above, we isolated DNA from the resulting population of cells and built sequencing libraries to quantify the representation of each single guide RNA. The raw output of all screens was reads per single guide RNA.

To select hits in our screens, we first normalized the output of our screens to reads per million, and then calculated the fold change in single guide RNA representation between different samples. For our priming screen, we focused on the fold change in single guide RNA representation between NGFR^HIGH^/EGFR^HIGH^ cells and the bulk population of melanoma cells. For the resistance screen, we focused on the fold change in single guide RNA representation between cells treated for three weeks with 1μM vemurafenib (a BRAF^V600E^ inhibitor) and cells never exposed to the drug. After normalizing the change in single guide RNA representation of each single guide RNA by the median change across all single guide RNAs, we organized our hits into tiers (one through four) based on the percent of single guide RNAs against the target exhibiting at least a two-fold change in representation. We considered high confidence hits those targets where (1) ≥ 75% (Tier 1) or ≥ 66% (Tier 2) of its single guide RNAs showed at least a two-fold enrichment/depletion throughout the screen, and (2) no two single guide RNAs showed a significant change (two-fold change) in opposing directions (i.e. one single guide RNA is significantly enriched in the selected population while another one is significantly depleted). Other targets that showed a two-fold enrichment/depletion throughout the screen, but in less than 66% of its single guide RNAs were considered lower confidence hits (Tier 3 and Tier 4). Note that we excluded from analysis any single guide RNA with less than 10 raw reads in all samples.

#### Secondary, targeted pooled CRISPR screen

To validate the replicability and generality of our hits, we designed a pool of single guide RNAs for targeted screening that targeted proteins that either emerged as hits in our initial screens or did not pass our hit-selection criteria but changed the frequency of NGFR^HIGH^/EGFR^HIGH^ cells or the frequency of cells resistant to vemurafenib (Supplemental Table 5). In this pool, we included ∼3 single guide RNAs per protein target, and carried out the screen in WM989-A6-G3-Cas9-5a3 cells as well as in another BRAF^V600E^ melanoma cell line, 451Lu-Cas9. As before, we conducted a priming screen where we isolated NGFR^HIGH^/EGFR^HIGH^ cells as well as a resistance screen where we exposed cells to 1μM vemurafenib for three weeks. Here too, we first normalized the output of our screens to reads per million, and then calculated the fold change in single guide RNA representation between different samples. Unlike on our initial screens, here we normalized the change in single guide RNA representation to the median change in representation of the ten non-targeting single guide RNAs controls included in the screen.

#### Tumor growth assays in patient-derived xenografts

All animal experiments were approved by the Institutional Animal Care and Use Committee (IACUC) (IACUC #112503X_0) and were performed in an Association for the Assessment and Accreditation of Laboratory Animal Care (AAALAC) accredited facility. WM989-A6-G3-Cas9-5a3 human melanoma cells (1 x 10^6^ cells) suspended in 100 ul of PBS were subcutaneously injected into 8-week-old NOD/SCID mice (Charles River Laboratories, Wilmington, MA). When resulting tumors reached 150 mm^3^, mice were fed either AIN-76A chow (untreated group, placebo) or AIN-76A chow containing 417 mg/kg PLX4720 (treated group). Tumor sizes were measured every 3-4 days using digital calipers, and tumor volumes were calculated using the following formula: volume = 0.5 x (length x width^2^). Mice were euthanized when tumors reached ∼1,500mm^3^ or upon development of skin necrosis.

To assess growth differences between knockouts and control tumors, we first quantified for each mouse the change in tumor size from the initial time point to the time point in question as a log2 fold change in tumor volume. We determined statistical significance of the differences observed between a knockout and controls at each therapy timepoint with a one-tailed t-test.

For each knockout cell line, we then calculated the mean tumor volume and standard error of the mean, which we report in Fig. 3. Note that within a given knockout-to-control comparison within each of the treatment arms we defined the endpoint as the last point in time at which at least 50% of the mice in each group (knockout and control cell line) were still alive.

#### Immunostains

For NGFR stain of fixed cells, after fixation and permeabilization, we washed the cells for 10 min with 0.1% BSA-PBS, and then stained the cells for 10 min with 1:500 anti-NGFR APC-labelled clone ME20.4 (Biolegend, 345107). After two final washes with PBS we kept the cells in PBS. For EGFR and NGFR stains of live cells, we incubated melanoma cells in suspension for 1 hour at 4C with 1:200 mouse anti-EGFR antibody, clone 225 (Millipore, MABF120) in 0.1% BSA PBS. We then washed twice with 0.1% PBS-BSA and then incubated for 30 minutes at 4C with 1:500 donkey anti-mouse IgG-Alexa Cy3 (Jackson Laboratories, 715-545-150). We washed the cells again (twice) with 0.1% BSA-PBA and incubated for 10 minutes with 1:500 anti-NGFR APC-labelled clone ME20.4 (Biolegend, 345107). We again washed the cells twice with 0.1% BSA-PBS and finally re-suspended them in 1%BSA-PBS.

#### Isolation of NGFR^HIGH^/EGFR^HIGH^ cells (MACS + FACS)

To enrich for NGFR^HIGH^/EGFR^HIGH^ cells we first immunostained melanoma cells as detailed above. Then, we used a Manual Separator for Magnetic Cell Isolation (MACS, with LS columns and Anti-APC microbeads). In short, following the manufacturer’s instructions, we incubated cells and microbeads at 4C for 15 min, then washed and pelleted the cells via centrifugation. After resuspending the cells, we passed them through LS magnetic columns. After enriching for NGFR^HIGH^ cells, we proceeded to select only the cells expressing both NGFR and EGFR via Fluorescent-Activated Cell Sorting (FACS, MoFlo Astrios EQ).

#### Growth of resistant colonies

To grow melanoma cells resistant to BRAF^V600E^ inhibition, we exposed melanoma cells to 1μM vemurafenib (PLX4032, Selleckchem S1267) for 2-3 weeks. For the BRAF^V600E^ and MEK co-inhibition assays, we also used dabrafenib at 500nM and 100nM (GSK2118436, Selleckchem S2807), trametinib at 5nM and 1nM (GSK1120212, Selleckchem S2673), and cobimetinib at 10nM and 1nM (GDC-0973, Selleckchem S8041).

#### Inhibition of DOT1L via small molecule inhibitor

For all assays involving pharmacological inhibition of DOT1L we used pinometostat at concentrations ranging from 1μM to 5μM (EPZ5676, Selleckchem S.7062).

#### MiSeq library construction and sequencing

In order to quantify the single guide RNA representation following selection in our screen we sequenced the single guide RNAs as per ^44^. In short, we isolated genomic DNA using the Quick-DNA Midiprep Plus Kit (Zymo Research: #D4075) per manufacturer specifications. We then PCR-amplified the single guide RNAs using Phusion Flash High Fidelity Master Mix Polymerase (Thermo Scientific: #F-548L) and primers that incorporate a barcode and a sequencing adaptor to the amplicon. Our amplification strategy consisted of an initial round of parallel PCRs (23-29 cycles of up to 200 parallel reactions per sample. We then pooled the PCR reactions and purified them using the NucleoSpin^®^ Gel and PCR Clean-up kit (Macherey-Nagel: #740609.250). We continued with eight PCR cycles using Phusion Flash High Fidelity Master Mix Polymerase, followed by column purification with the QIAquick PCR Purification Kit (QIAGEN: #28106). We quantified the single guide RNA libraries with the DNA 1000 Kit (Agilent: #5067-1504) on a 2100 Bioanalyzer Instrument (Agilent: #G2939BA). We pooled the barcoded single guide RNA libraries and sequenced via 150-cycle paired-end sequencing (MiSeq Reagent Kit v3, Illumina: #MS-102-3001). We then mapped the resulting sequences to our reference single guide RNA library and proceeded to select hits.

#### Cell fixation and permeabilization

For our imaging assays we fixed cells for 10 min with 4% formaldehyde and permeabilized them with 70% ethanol overnight.

#### Colony formation assays

For each condition tested, we first plated cells in duplicate (∼10-50,000 cells per well of a 6-well plate). We fixed and permeabilized one of the duplicates to use as a baseline and exposed the second duplicate to the test condition. At the endpoint, we fixed and permeabilized the second duplicate.

#### Image analysis of NGFR immunostains

We developed a custom MATLAB pipeline for counting cells and quantifying immunofluorescence signal of DAPI-stained and NGFR-stained cells (https://bitbucket.org/arjunrajlaboratory/rajlabscreentools/src/default/). The software stitches together a large tiled image, then uses DAPI to identify cells. Using the nuclear area, it then looks at a set of pixels near the nucleus to quantify fluorescence intensity of the NGFR staining. After quantifying the expression level of NGFR following knockout of select screen targets and of non-targeting controls, we quantified the minimum expression level needed to be considered an NGFR^HIGH^ cell. First, we selected the top one percent highest expressors of NGFR in each of our non-targeting negative controls. Then, within that top one percent we obtained the median expression level of the lowest expressor across all controls, and used that as a threshold to quantify the frequency of NGFR^HIGH^ cells in each of our knockout samples. Then, we calculated the change in frequency of NGFR^HIGH^ cells in each test condition compared to controls and obtained a median fold change and standard deviation across all samples with knockout of one same protein (∼3 different biological samples per protein). In total, we targeted ∼86 different proteins across ∼258 different knockout biological samples.

#### Image analysis of colony formation

We developed a custom MATLAB pipeline for counting cells and colonies in tiled images of DAPI-stained cells (https://bitbucket.org/arjunrajlaboratory/colonycounting_v2/src/default/). First, the software stitches the individual image tiles into one large image by automatically (or with user input) determining the amount of overlap between each individual image. Then, the software identifies the location of each cell in the stitched image by searching for local maxima. We then manually identify the colony boundaries and quantify the number of colonies in each sample. We then calculate the frequency of resistant colonies by dividing the number of colonies by the total number of cells present in culture prior to BRAF^V600E^ inhibition. Finally, we scale the frequency of colonies to colonies per 10,000 cells and calculate the change in frequency between each sample and the median change across controls.

#### RNA-sequencing and identification of differential expression

We sequenced mRNA in bulk from WM989-A6-G3 and WM989-A6-G3-Cas9 populations as per Shaffer et. al. In addition to quantifying the transcriptome of EGFR^HIGH^cells, NGFR^HIGH^, NGFR^HIGH^/EGFR^HIGH^ cells and vemurafenib-resistant cells, we quantified the transcriptional changes following the knockout of many tier 1 and tier 2 hits from both the priming and resistance screens. In addition to hits from our screens, we also quantified the transcriptome of targets that were not tier 1 or tier 2 hits, but show a change in the frequency of NGFR^HIGH^/EGFR^HIGH^ cells or of cells resistant to vemurafenib. In total, we targeted ∼83 different proteins, each in triplicate (using different single guide RNAs) for a total of 280+ RNA sequencing samples. For each sample, we isolated mRNA and built sequencing libraries using the NEBNext Poly(A) mRNA Magnetic Isolation Module and NEBNext Ultra RNA Library Prep Kit for Illumina per manufacturer instructions. We then sequenced the libraries via paired-end sequencing (36×2 cycles) on a NextSeq 500. We aligned reads to hg19 and quantified reads per gene using STAR and HTSeq. We then used DEseq2 to identify differentially expressed genes.

#### Gene set enrichment analysis

To identify “biological signatures” enriched or depleted following the knockout of a given target we used the GSEA software (http://software.broadinstitute.org/gsea/index.jsp). We focused in the Biological Process ontology of the Gene Ontology gene sets (http://geneontology.org) to obtain enrichment scores.

#### Grouping of targets based on transcriptomic analysis

To group targets into classes based on their transcriptional effects, we clustered all RNA-seq samples (hierarchical clustering via pheatmap in R) based on the change in expression (as obtained by DEseq2) of any gene differentially expressed (two-fold change over control, with an adjusted p value ≤ 0.05) in at least one of the 83+ knockouts. We also grouped targets via pheatmap based on the enrichment scores obtained via GSEA. To identify the axes that account for the variability between each knockout we also performed principal component analysis based on the gene set enrichment scores of each knockout. Note that in the aforementioned analysis we included the transcriptomes of pre-resistant cells (marked by the expression of EGFR alone, NGFR alone, and NGFR and EGFR in combination) and of cells resistant to vemurafenib.

#### Software and data availability

All data and code used for the analysis can be found at https://www.dropbox.com/sh/t08558cl4mepfm6/AABBvbtlTPSNNPoMC9NTro-9a?dl=0 The software used for colony growth image analysis can be found at: https://bitbucket.org/arjunrajlaboratory/colonycounting_v2/src/default/. The software used for analysis of immunofluorescence images can be found at: https://bitbucket.org/arjunrajlaboratory/rajlabscreentools/src/default/

## Supplemental Figure legends

**Supplemental Figure 1.**
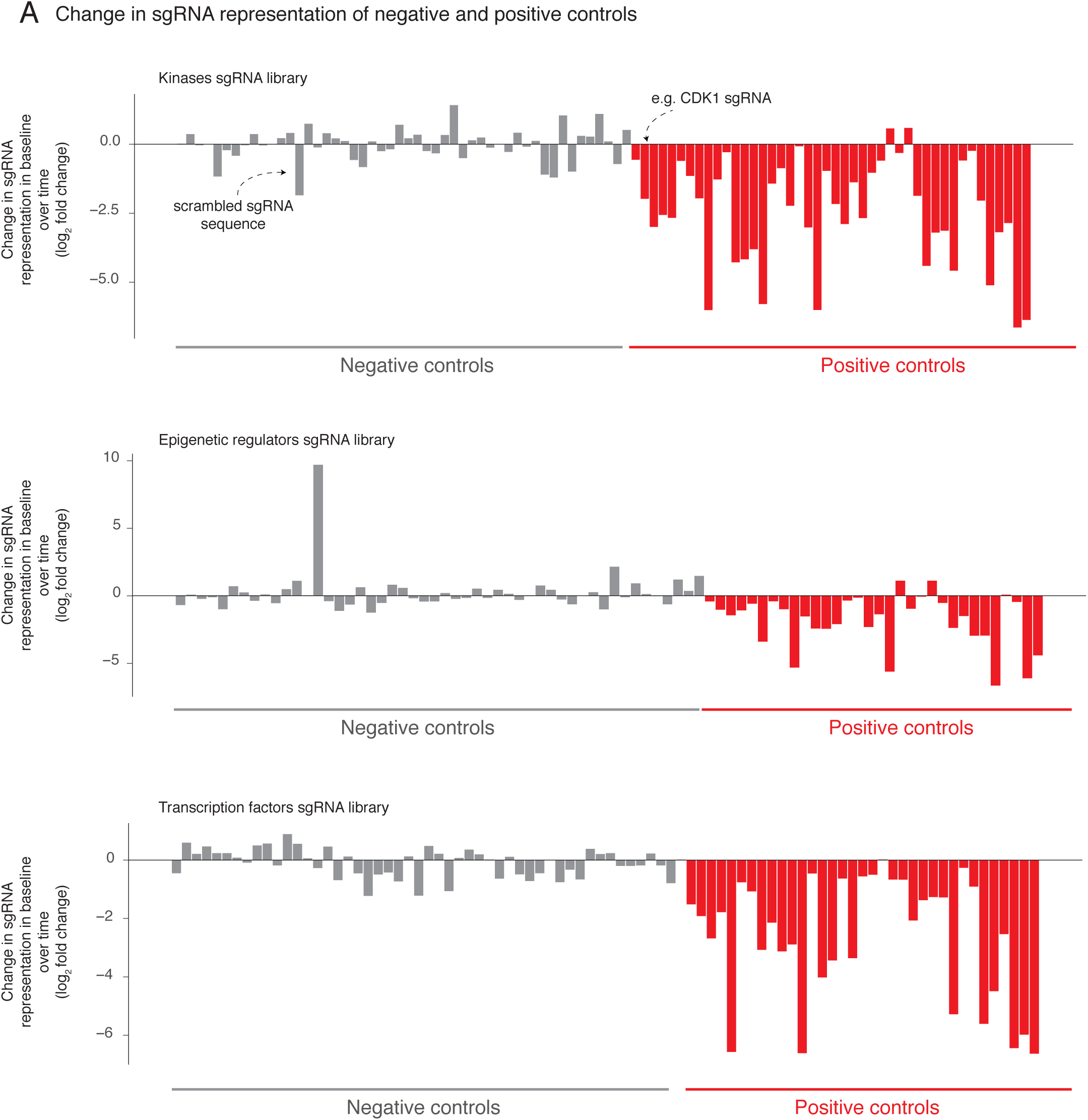
Effect of negative and positive control single guide RNAs in the CRISPR screens. Our pooled CRISPR screen included non-targeting single guide RNAs as negative controls (gray bars, 50+ single guide RNAs) as well as single guide RNAs affecting cell viability as positive controls (red bars, 25+ single guide RNAs). We quantified the change in representation of these single guide RNAs over time and report the log_2_ fold change in representation from six days after transfection to right before selection (vemurafenib exposure or selection by NGFR and EGFR expression). We expect positive controls to lose representation over time more often than negative controls. Our screening scheme utilized three separate pooled single guide RNA libraries, one targeting kinases (top), another targeting epigenetic domains (middle), and a final one targeting transcription factors (bottom).

**Supplemental Figure 2.**
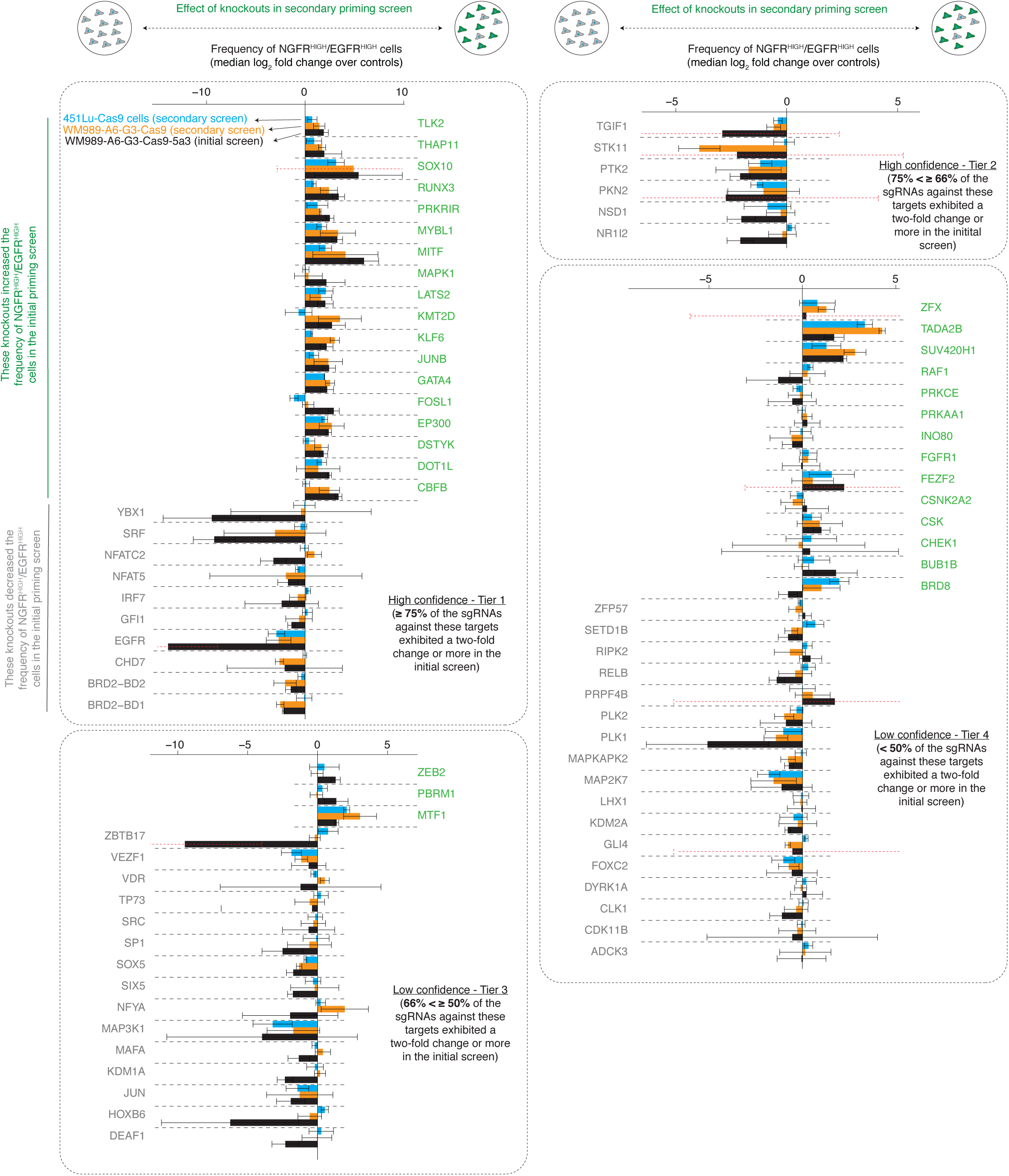
Secondary validation of hits across multiple cell lines by secondary targeted CRISPR screening. We assessed the robustness and generality of the effect of hits identified in the priming screen (WM989-A6-G3-Cas9-5a3, black bars) by carrying out a secondary screen containing single guide RNAs for 34 of the high confidence hits (Tiers 1 and 2) we identified in the priming screen, as well as another 52 lower confidence factors from Tiers 3 and 4 (these lower confidence hits may also have been high confidence hits in the resistance screen). See Supplemental Table 5 for a list of the targets. We carried out this screen in WM989-A6-G3-Cas9 (orange bars) as well as in another BRAF^V600E^ melanoma cell line (451Lu-Cas9, blue). Within each tier, names labeled in green correspond to targets whose single guide RNAs are enriched in NGFR^HIGH^/EGFR^HIGH^ cells in the initial screen and gray represents targets whose single guide RNAs are underrepresented in these rare cells. We plotted the median log_2_ fold change in single guide RNA representation (normalized by non-targeting controls) across three single guide RNAs. Error bars represent the standard deviation of the fold change across all single guide RNAs for a given target. Dotted error bars in red extend beyond the limits of the graph. Note that the limits of the axes vary between tiers. We found that 25 of the 34 high confidence hits showed at least a two fold change in the frequency of NGFR^HIGH^/EGFR^HIGH^ cells concordant with the effects detected in the original screening clonal cell line (WM989-A6-G3-Cas9-5a3). In 451Lu-Cas9 cells, 20 of the 34 targets also showed a change in the frequency of NGFR^HIGH^/EGFR^HIGH^ cells, with 11 of those exhibiting at least a two-fold change. Although quantitatively they were not as strong as the effects of the Tier 1 and 2 hits, even Tier 3 and 4 hits displayed qualitative agreement in these secondary screens.

**Supplemental Figure 3.**
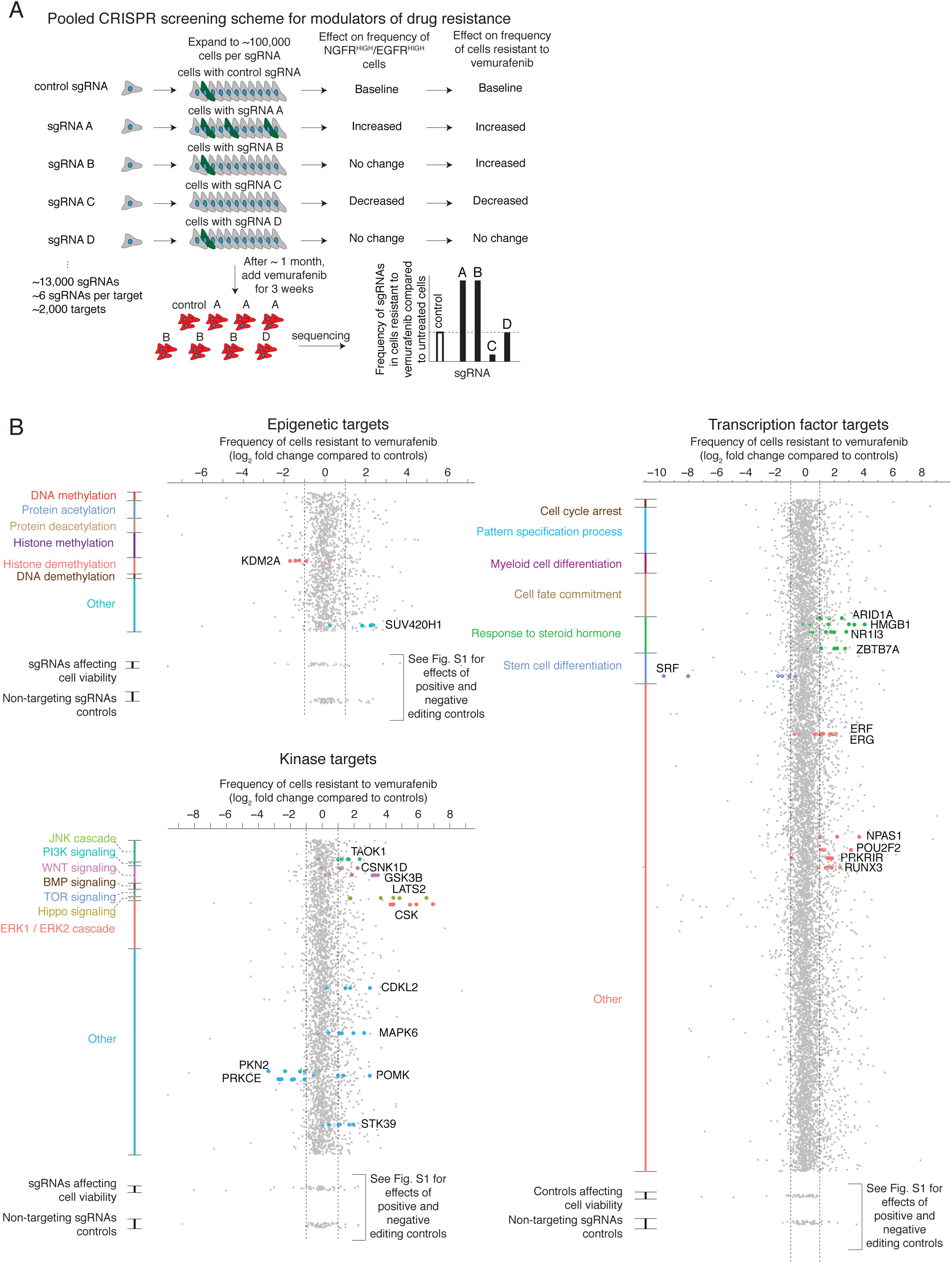
Screen for factors modulating number of resistant colonies upon BRAF^V600E^ inhibition. **A.** We performed a pooled CRISPR screen to detect modulators of the number of drug-resistant cells that grow in the presence of the BRAF^V600E^ inhibitor vemurafenib. After transducing a library of single guide RNAs and expanding the population, we exposed the cells to the BRAF^V600E^ inhibitor vemurafenib (1µM) for 3 weeks, after which we sequenced the single guide RNAs in the surviving population. Changes in the frequency of detection of a given single guide RNA indicates that its target may affect the ability of a cell to survive and proliferate upon BRAF^V600E^ inhibition. **B.** After transfecting a population of melanoma cells, we exposed them to vemurafenib (BRAF^V600E^ inhibitor, 1μM) for three weeks to grow resistant colonies. We then sequenced the DNA to quantify the single guide RNA representation of each target in the resulting population, using the same libraries as in Fig. 1. As before, we ranked the targets into tiers based on the percent of single guide RNAs that exhibited at least a two-fold change in representation throughout the screen (Tier 1, ≥ 75%; Tier 2, ≥ 66%; Tier 3, ≥ 50%; Tier 4, < 50%), thus reflecting the degree of confidence we have in the hit (High confidence hits: Tiers 1 and 2; Low confidence hits: Tiers 3 and 4). In this screen, we identified 24 high confidence factors. For a more detailed description see the methods section.

**Supplemental Figure 4.**
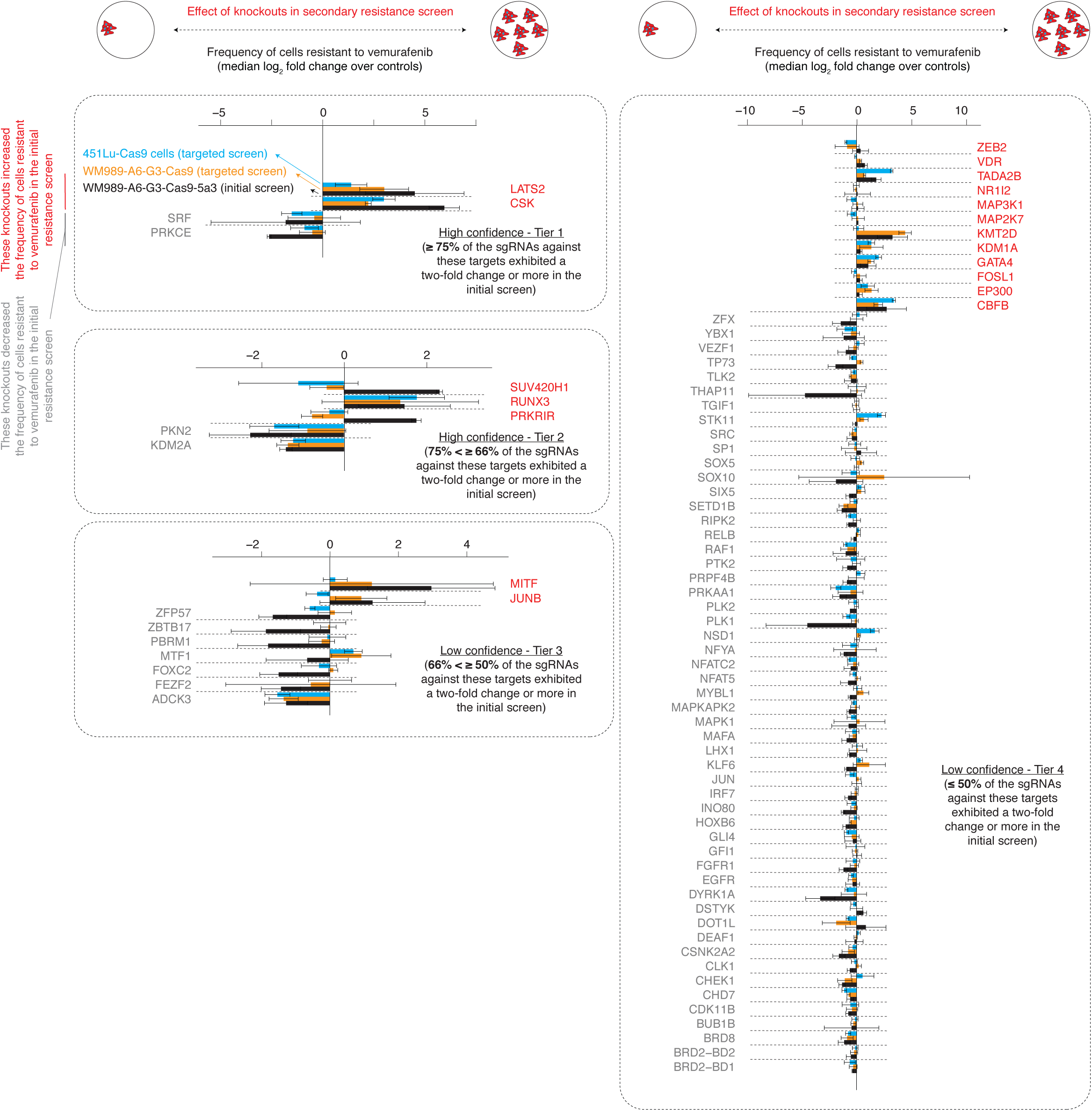
Secondary validation of hits across multiple cell lines by secondary targeted CRISPR screening. We assessed the robustness and generality of the effect of hits identified in the initial resistance screen (WM989-A6-G3-Cas9-5a3, black bars) by carrying out a secondary screen containing single guide RNAs for nine high confidence targets (as well as 77 targets that either affected vemurafenib resistance but did not pass the thresholds to be called a hit, or affected the frequency of NGFR^HIGH^/EGFR^HIGH^ cells; Supplemental Table 5 for a list of all the targets). We carried out this screen in WM989-A6-G3-Cas9 (orange bars) as well as in another BRAF^V600E^ melanoma cell line (451Lu-Cas9, blue). Within each tier, names labeled in red correspond to targets whose single guide RNAs are enriched in cells resistant to vemurafenib in the initial screen and gray represents targets whose single guide RNAs are underrepresented in these cells. We plotted the median log_2_ fold change in single guide RNA representation (normalized by non-targeting controls) across three single guide RNAs. Error bars represent the standard deviation of the fold change across all single guide RNAs for a given target. Dotted error bars in red extend beyond the limits of the graph. Note that the limits of the axes vary between tiers. In WM989-A6-G3-Cas9, we found that seven of the nine targets replicated the effect that we observed originally. For 451Lu-Cas9, the same seven factors showed similar effects.

**Supplemental Figure 5.**
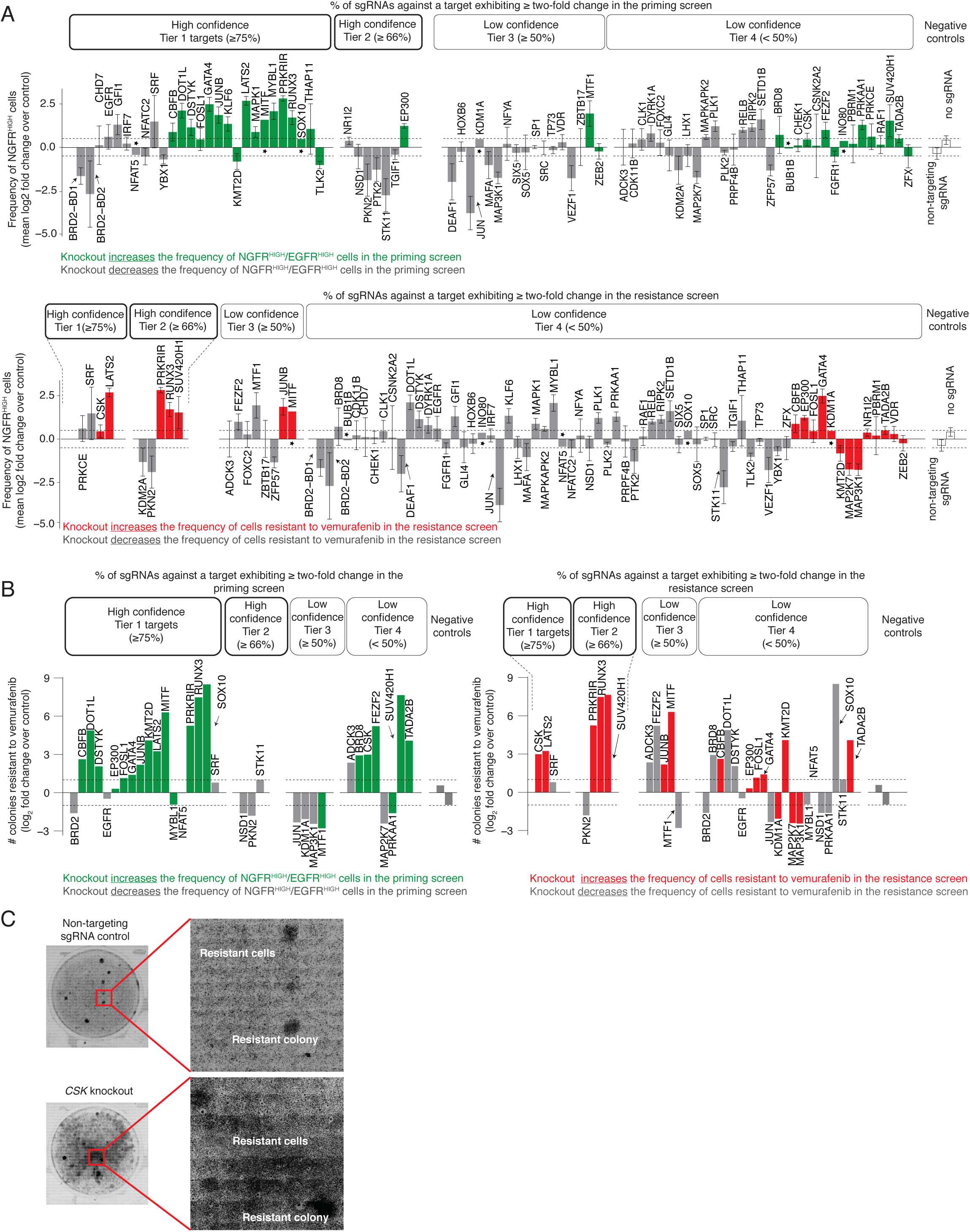
Validation of effects of hits from priming and resistance screens by via NGFR immunofluorescence and resistant colony formation. **A.** Frequency of NGFR^HIGH^ cells following the knockout of select targets. Each bar represents the change in the number of NGFR^HIGH^ cells following knockout of the gene indicated over replicates, each using a different single guide RNA. Error bars represent the standard error of the mean across the replicates. We carried out each measurement over three replicates, but excluded samples with low cell density (< 500 cells). The star above or below the bars indicate targets where, after excluding samples with low cell numbers, n = 1. Tier refers to the degree of confidence we have in each particular hit, with tier 1 representing highest confidence hits for which ≥ 75% of the single guide RNAs passed a threshold of two-fold change in the initial screens. We performed this analysis for hits from both the priming screen (top) and the resistance screen (bottom). 21 of 34 high confidence hits from the priming screen showed at least a 50% increase or decrease in the frequency of NGFR^HIGH^ cells over control. Of the lower confidence hits (Tier 3 and Tier 4) 21 out of 49 targets increased or decreased the frequency of NGFR^HIGH^ cells by 50% or more. **B.** Resistance phenotype of melanoma cells following the knockout of hits from the initital screens. Each bar represents the log_2_ fold change over non-targeting control in the number colonies able to grow in vemurafenib following knockout of the gene indicated. The number of colonies for each target is normalized to the number of cells present in culture before BRAF^V600E^ inhibition, reported as number of colonies per every 10,000 cells in culture prior to treatment (see methods). As before, the different tiers represent the percent of single guide RNAs against a given target exhibiting at least a two-fold change throughout the initial (left) priming or (right) resistance screens. On the left panel, we labeled in green and gray the effect a given target has in the frequency of NGFR^HIGH^/EGFR^HIGH^ cells (based on the initial priming screen). On the right panel, we labeled in red and gray the effect a given target has in the number of cells that resist BRAF^V600E^ inhibition (based on the initial resistance screen). In this plot each bar represents one experimental replicate. See Supplemental Fig. 6 for replicates. **C.** These images show the effect of CSK knockout on a cell’s ability to develop resistance to BRAF^V600E^ inhibition. Here, we plated CSK-knockout WM989-A6-G3-Cas9-5a3 cells and exposed them to 1μM vemurafenib for three weeks. As before, after the treatment we counted the number of resulting colonies and compared it to the number of colonies resulting from WM989-A6-G3-Cas9-5a3 cells without the knockout. Note that the number of resistant cells in the CSK sample is too large to accurately identify individual colonies. We only counted colonies we could clearly delineate, and thus, the number of colonies reported is an underestimate.

**Supplemental Figure 6.**
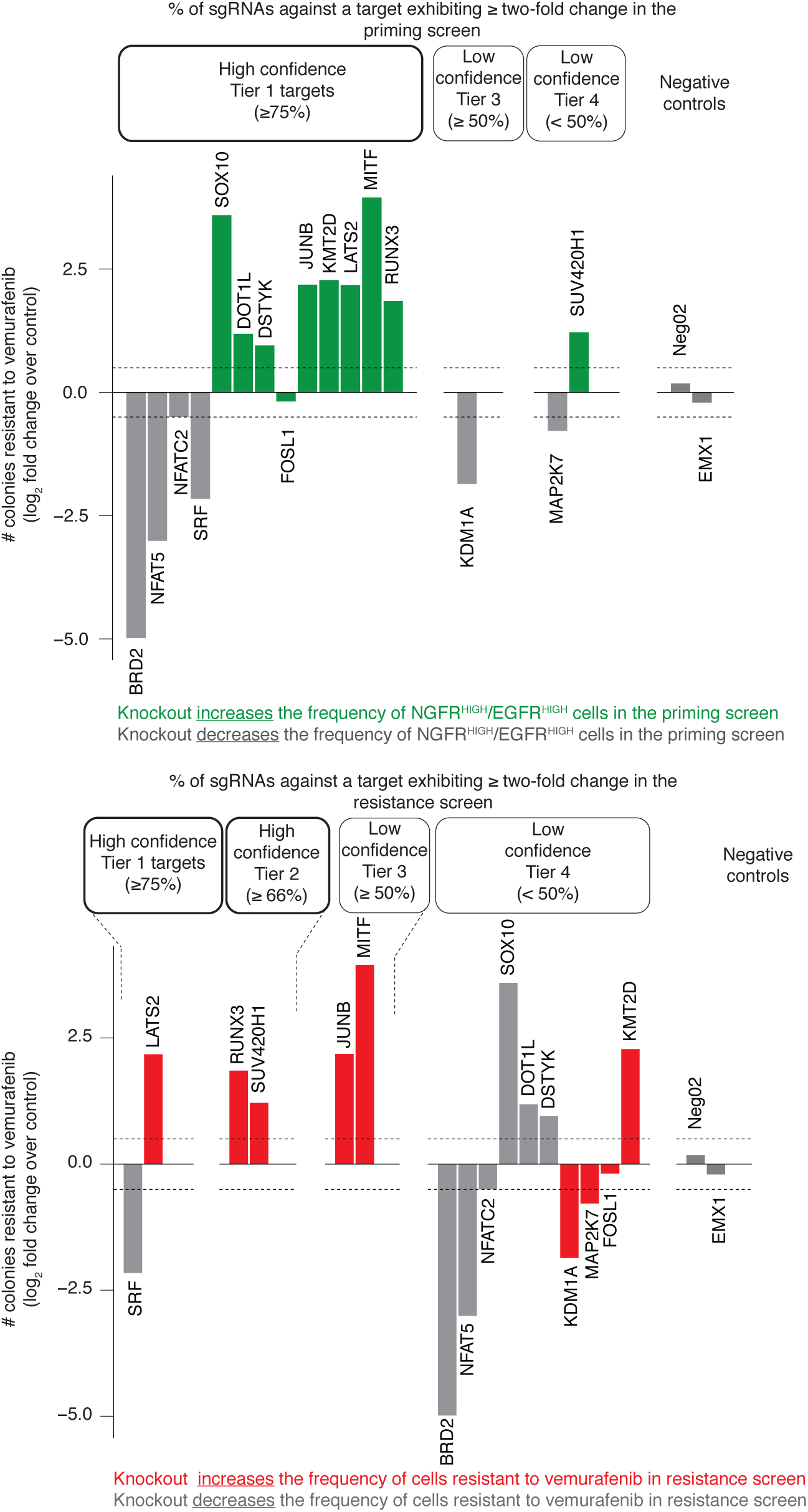
Validation of effects hits by resistant colony formation. Resistance phenotype of melanoma cells following the knockout of hits from the initial screens. Each bar represents the log_2_ fold change over non-targeting control in the number of colonies able to grow following knockout of the gene indicated. The number of colonies for each target is normalized to the number of cells present in culture before BRAF^V600E^ inhibition, reported as number of colonies per every 10,000 pre-treatment cells (see methods). As before, the different tiers represent the percent of single guide RNAs against a given target exhibiting at least a two-fold change throughout the initial (top) priming or (bottom) resistance screens. On the top panel, we labeled in green and gray the effect a given target has in the frequency of NGFR^HIGH^/EGFR^HIGH^ cells (based on the initial priming screen). On the bottom panel, we labeled in red and gray the effect a given target has in the number of cells that resist BRAF^V600E^ inhibition (based on the resistance screen). In this plot each bar represents one experimental replicate (distinct from the one in Supplemental Fig. 5B).

**Supplemental Figure 7.**
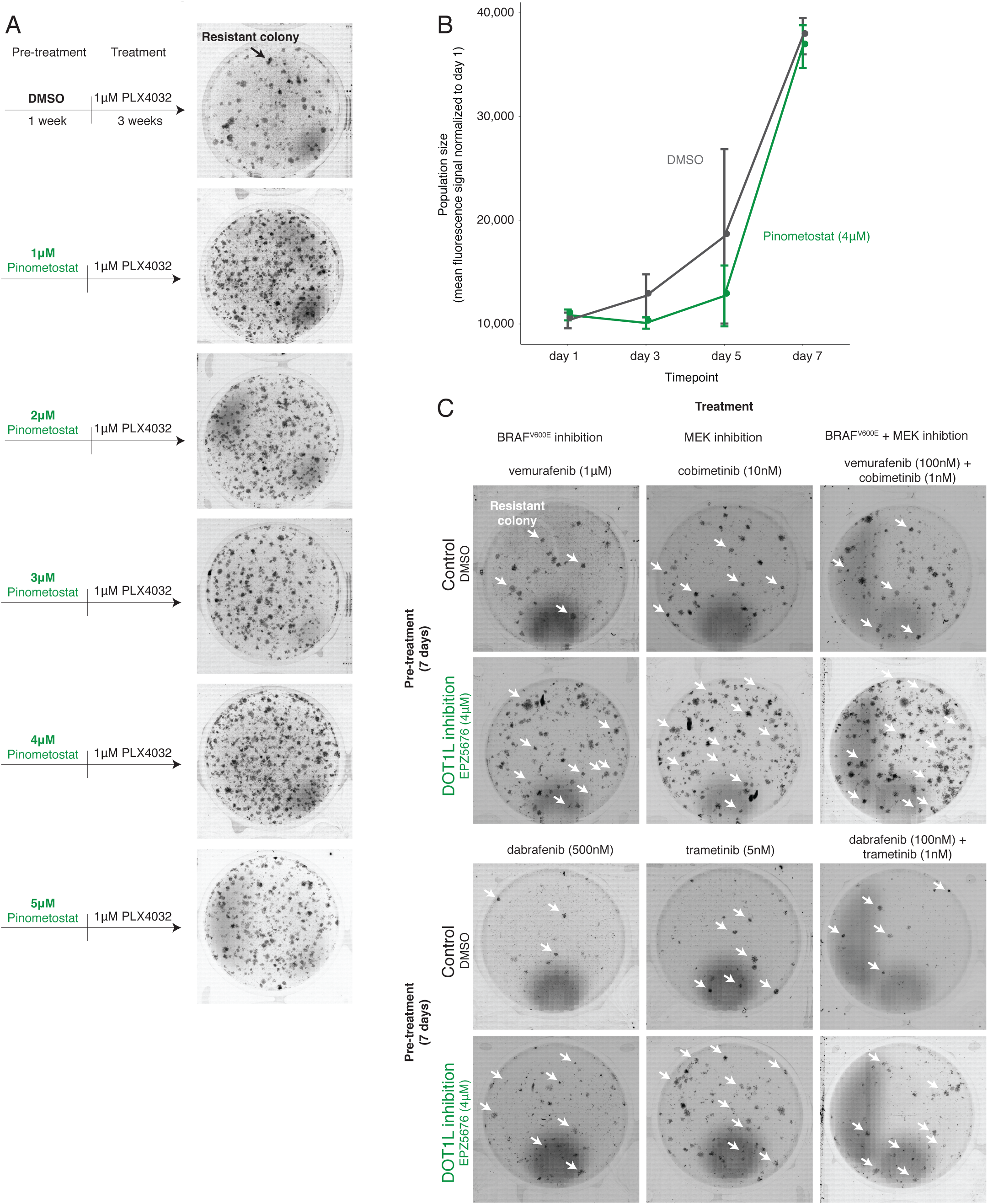
Effect of pharmacological inhibition of DOT1L on resistance to BRAF^V600E^ and MEK inhibition. **A.** Resistance phenotype of melanoma cells following pharmacological inhibition of DOT1L. We pre-treated melanoma cells for seven days with either DMSO, or various concentrations of the DOT1L inhibitor pinometostat (EPZ5676). Then, we exposed the cells to 1μM vemurafenib for three weeks. **B.** To assess the effect of DOT1L inhibition on cellular proliferation, we the compared the population size of WM989-A6-G3 cells over time treated with either 4μM of pinometostat (DOT1L inhibitor) or with DMSO. The population size is estimated by the amount of nucleic acids present in the population using a CyQuant GR dye. The values represent mean fluorescence signal over triplicates. Error bars represent standard error of the mean. **C.** Resistance phenotype of melanoma cells to BRAF^V600E^ and MEK inhibitors following pharmacological inhibition of DOT1L. We pre-treated melanoma cells for seven days with either DMSO or 4μM of pinometostat. We then exposed the cells to one of two BRAF^V600E^ inhibitors (vemurafenib and dabrafenib, left panels), to one of two MEK inhibitors (cobimetinib and trametinib, middle panels), or to a combination of a BRAF^V600E^ and MEK inhibitor (vemurafenib + cobimetinib; dabrafenib + trametinib, right panels). White arrows point to a few of the many colonies that grew under each condition.

**Supplemental Figure 8.**
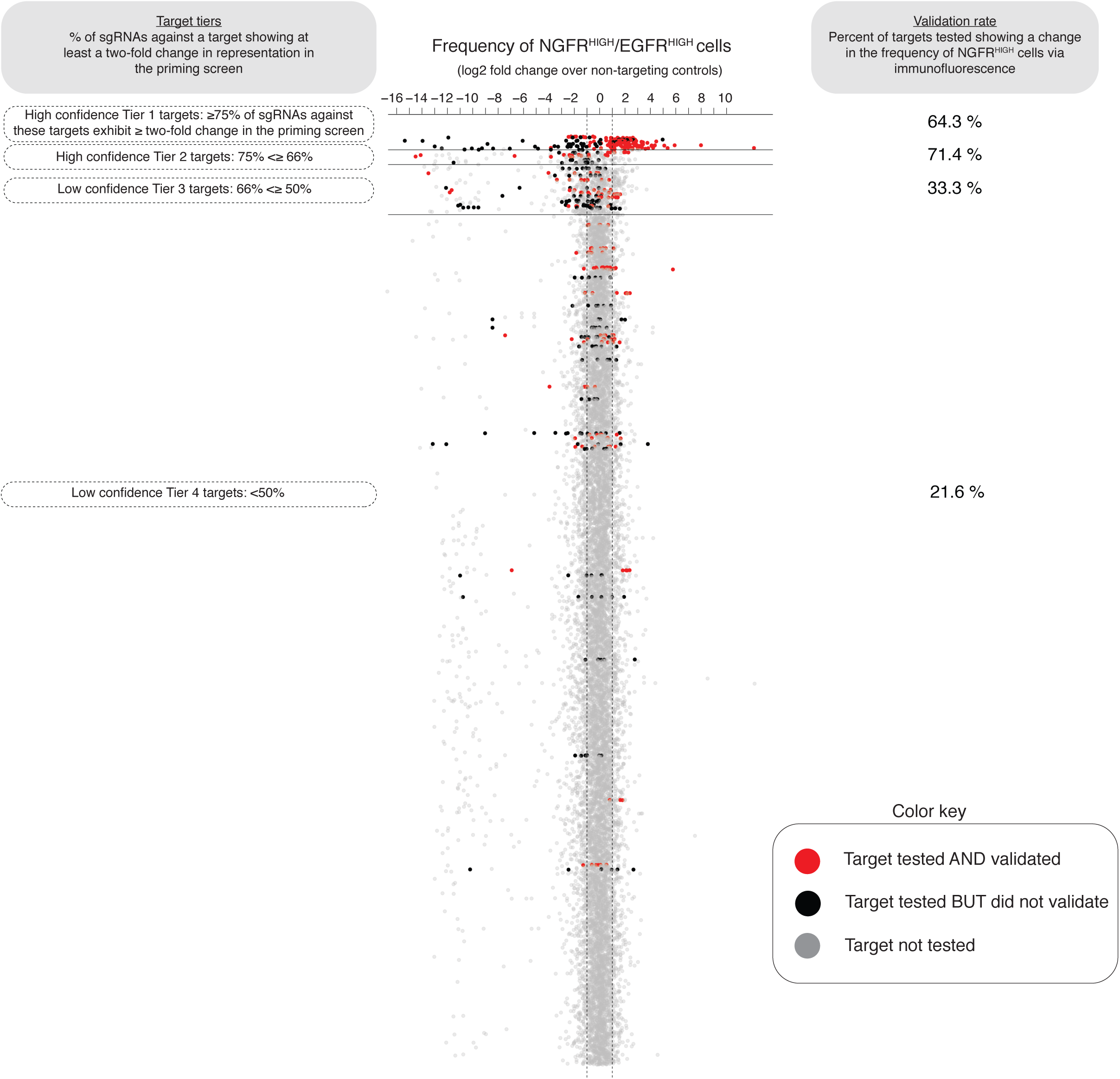
Percent of targets from the priming screen that validate. To assess the sensitivity of our screen, we validated the effect observed in the initial priming screen for a select group of targets via NGFR immunofluorescence. Here, each dot represents an individual single guide RNA, and we plot the change in single guide RNA representation between NGFR^HIGH^/EGFR^HIGH^ cells and controls (as measured in the priming screen). We then organize all sgRNAs into tiers (y-axis, tiers one through four) based on the percent of single guide RNAs against a target showing at least a two-fold change in representation on NGFR^HIGH^/EGFR^HIGH^ cells. In red, we labeled targets that when tested again produced at least a 50% change in the frequency of NGFR^HIGH^ cells. In black, we labeled targets that we tested but did not validate, and in gray we show targets we did not test. We display the percent of genes tested and validated at each tier on the right.

**Supplemental Figure 9.**
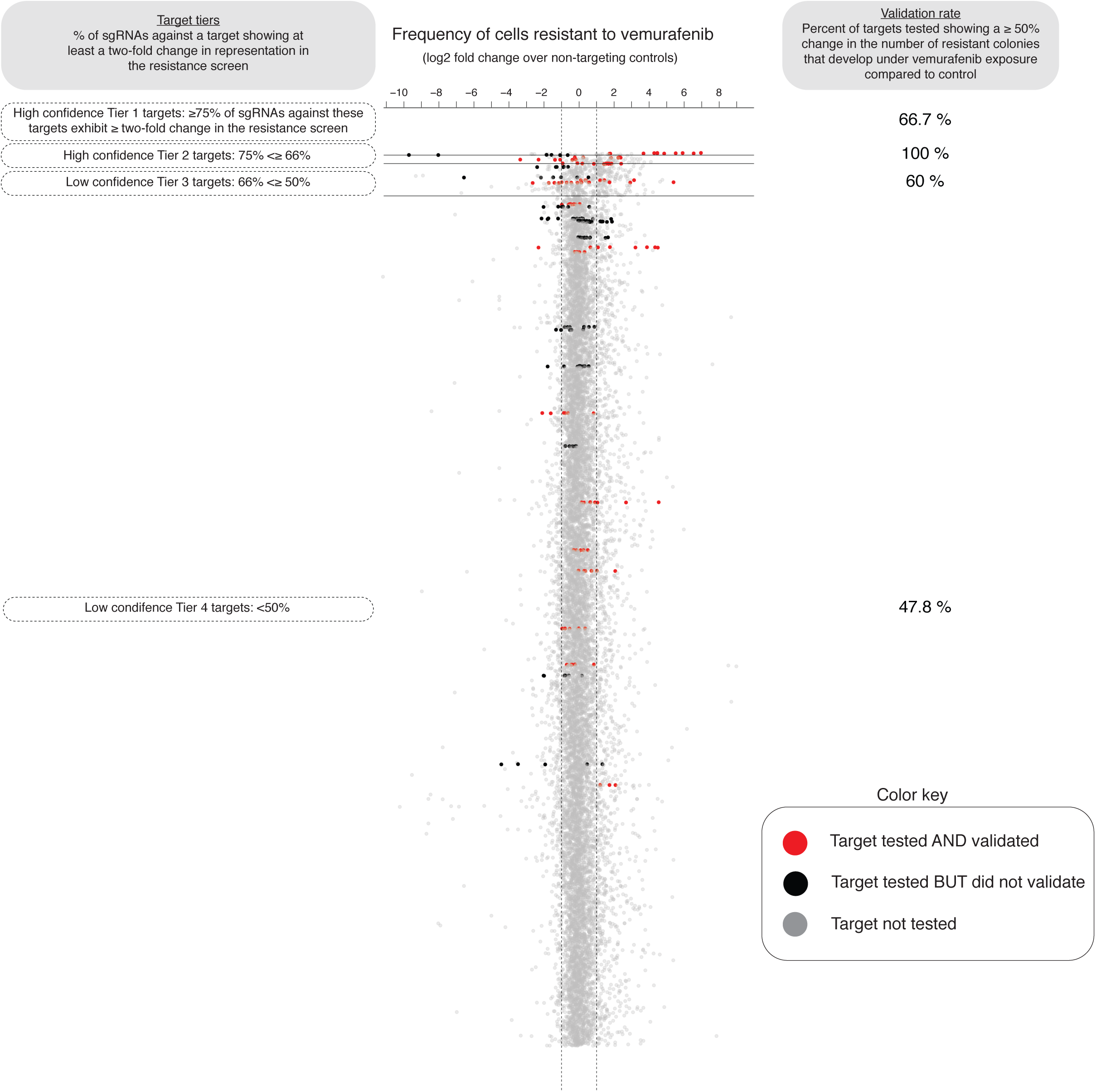
Percent of targets from the resistance screen that validate. To assess the sensitivity of our screen, we validated the effect observed in the initial resistance screen for a select group of targets via colony formation assays. Here, each dot represents an individual single guide RNA, and we plot the change in single guide RNA representation between cells resistant to vemurafenib and cells that have never been exposed to the drug (as measured in the resistance screen). We then organize all single guide RNAs into tiers (y-axis, tiers one through four) based on the percent of single guide RNAs against a target showing at least a two-fold change in representation on drug resistant cells. In red, we labeled targets that when tested again produced at least a 50% change in the frequency colonies resistant to BRAF^V600E^ inhibition. In black, we labeled targets that we tested but did not validate, and in gray we show targets we did not test. We display the percent of genes tested and validated at each tier on the right.

**Supplemental Figure 10.**
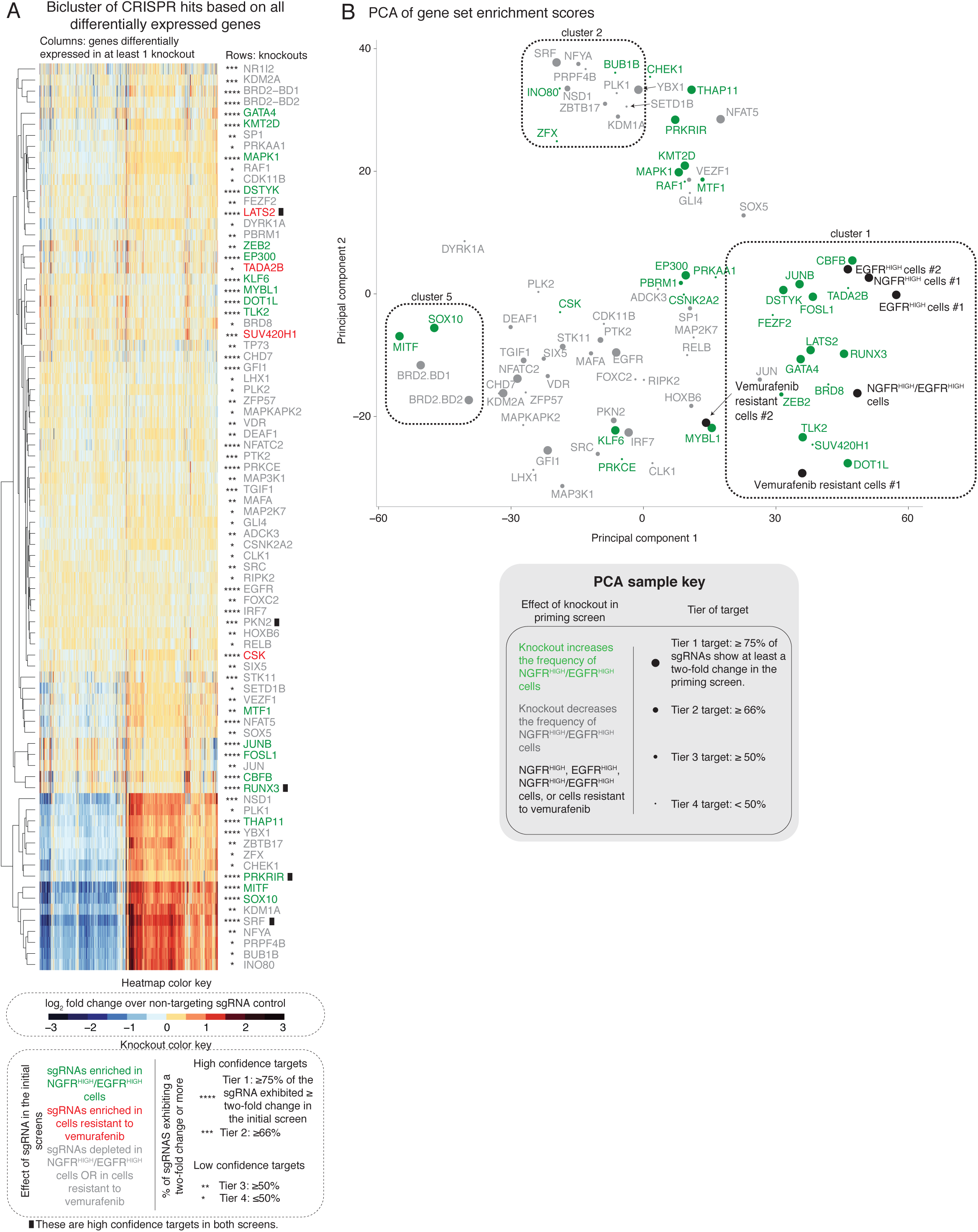
Transcriptional effects induced by knockout of select screen targets. **A.** The heatmap represents the biclustering analysis of different screen targets (rows) based on the change in expression of all genes differentially expressed in at least one knockout (columns). Within the heatmap, red indicates an increase in expression following the knockout, while blue indicates a decrease in gene expression (see heatmap color key). Each target (rows) represents transcriptomes of biological triplicates (unless otherwise stated on Supplemental Table 4). Target labels (rows) in green indicate genes whose knockout increased the frequency of NGFR^HIGH^/EGFR^HIGH^ cells in the initial screen. In red are those whose knockout increased the number of cells resistant to vemurafenib, and in gray are those that decreased the frequency of either NGFR^HIGH^/EGFR^HIGH^ cell or of cells resistant to vemurafenib. As before, we organized targets into confidence tiers indicated by the number of asterisks, based on the percent of single guide RNAs against that target that showed an effect in the initial screen (see knockout color key). **B.** We performed principal component analysis of the transcriptional effects induced by the knockout of select screen targets. We used as input the gene set enrichment scores from Fig. 5A to identify primary axes that account for the greatest degree of transcriptome variability across knockout cell lines. The color indicates the effect of the knockout in the initial priming screen. The size of the dot indicates the degree of confidence we have in each particular hit based on the percent of the single guide RNAs against a target that passed a threshold of two-fold change in the initial priming screen. In black, we labeled melanoma cells where we did not knockout any targets but either enriched for EGFR^HIGH^ cells, NGFR^HIGH^ cells, EGFR^HIGH^/NGFR^HIGH^ cells, or for cells resistant to vemurafenib.

**Supplemental Figure 11.**
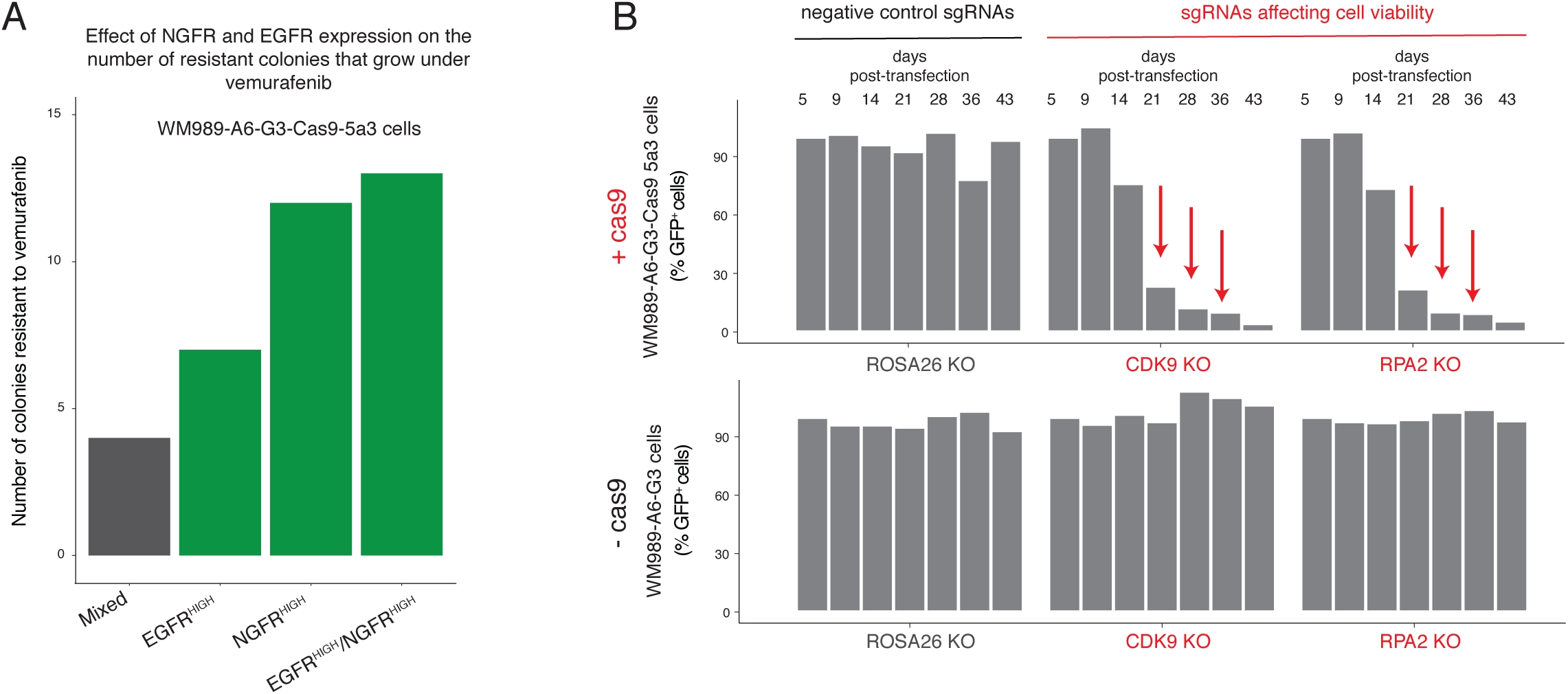
Technical validation of WM989-A6-G3-Cas9-5a3 cell line. **A.** WM989-A6-G3-Cas9-5a3 cells expressing NGFR, EGFR, and both NGFR and EGFR are more likely to survive and proliferate in the presence of vemurafenib ^14^. Here, we show the number of colonies that grow upon vemurafenib exposure in a mixed population of WM989-A6-G3-Cas9-5a3 or in the same population but enriched for EGFR^HIGH^ cells, NGFR^HIGH^ cells, or NGFR^HIGH^/EGFR^HIGH^ cells. **B.** In this plot, we show the single guide RNA representation (as percent GFP-positive cells) of controls over time in WM989-A6-G3 cells with or without Cas9 expression. Negative controls (black) are single guide RNAs aimed at ROSA26, a non-expressing gene in human melanoma. Positive controls (red) target proteins necessary for cell viability. Only cells expressing both Cas9 and a positive control single guide RNA should disappear from the population over time.

## Supplemental Tables

All tables can be found at: https://www.dropbox.com/sh/t08558cl4mepfm6/AABBvbtlTPSNNPoMC9NTro-9a?dl=0

